# Benchmarking Bayesian colocalization methods in validating Mendelian randomization-identified targets

**DOI:** 10.1101/2024.10.14.617627

**Authors:** Wenmin Zhang, Satoshi Yoshiji, Robert Sladek, Josée Dupuis, Tianyuan Lu

## Abstract

Mendelian randomization (MR) is an important tool for identifying potential biomarkers and drug targets. Colocalization analysis is crucial for validating MR findings and guarding against confounding due to linkage disequilibrium. We aim to benchmark the performance of four Bayesian colocalization methods in validating MR-based target discoveries from circulating proteins for cardiometabolic traits. We assessed the associations between circulating levels of 1,535 proteins and five cardiometabolic traits, followed by colocalization analyses using coloc, coloc+SuSiE, PWCoCo and SharePro. All methods demonstrated well-controlled false discoveries. SharePro demonstrated the highest frequency in supporting 160 (79.6%) of the 201 Bonferroni-significant protein-trait associations identified by MR, compared to coloc (supporting 40.3% of these associations), coloc+SuSiE (46.8%), and PWCoCo (45.8%), and was robust to varying prior colocalization probabilities. Protein-trait associations supported by SharePro were more likely to agree with significant gene-level associations identified in exome-wide association studies and implicate known drug targets. Eight protein-trait associations were exclusively supported by SharePro, suggesting potential cardiometabolic biomarkers or drug targets, such as HSF1 and HAVCR2. In summary, SharePro most often supports statistically significant associations identified through MR for cardiometabolic traits. Combining multiple lines of evidence using different methods may substantially increase the yield of biomarker and drug target discovery programs.

## Introduction

The rapidly increasing burden of cardiometabolic diseases, including ischemic heart diseases, stroke, and type 2 diabetes, is presenting significant challenges for healthcare systems globally[1,2]. Identifying novel biomarkers and drug targets tailored to the molecular pathways underlying these diseases is crucial for improving healthcare outcomes[3]. However, due to the multifactorial nature of cardiometabolic diseases, the identification of effective biomarkers and drug targets remains challenging, costly and time-consuming[4].

Circulating proteins are attractive candidates for biomarker and drug target discovery, as they often have important roles as messengers, regulators, and effectors of physiological and pathological processes[5–7]. Recent technological advances have significantly enhanced our ability to measure and modify the circulating levels of proteins. However, unraveling the potential causal roles of circulating proteins in cardiometabolic diseases, for example, in randomized controlled trials, is challenging, given the high cost, ethical concerns, and logistical challenges. Traditional observational studies, while informative in identifying associations, can be susceptible to biases introduced by confounding and reverse causation.

In the past decade, Mendelian randomization (MR) has emerged as an important tool for testing potential causal effects while mitigating biases from confounding and reverse causation[8–10]. MR utilizes genetic variants as instruments for a given exposure, such as the circulating level of a specific protein, to assess the potential causal effect of the exposure on an outcome. Three essential assumptions are required in selecting instrumental variables for MR analyses. First, the genetic instrument should be a strong predictor of the exposure; second, the genetic instrument should not be associated with any confounders; third, there is no horizontal pleiotropy, where the genetic instrument should affect the outcome through the specified exposure and not through any other mechanisms.

Lead variants of cis-protein quantitative trait loci (pQTLs), characterized in recent genome-wide association studies (GWAS) of circulating proteins[5–7], can serve as genetic instruments for MR analyses. These variants are significantly associated with the circulating protein levels, have a reduced risk of confounding due to randomization at conception, and are likely to directly influence circulating levels of the associated proteins due to their proximity to the protein-coding genes. However, another important source of bias in MR is confounding due to linkage disequilibrium (LD)[11]. Specifically, if two genetic variants in LD independently influence the exposure and the outcome through distinct mechanisms, the instrumental variable assumption of MR would be violated. To guard against this potential bias, colocalization analyses have been employed to evaluate whether the exposure and the outcome share the same causal variants[11].

One of the first and most commonly used methods for colocalization analyses is coloc[12]. For a pair of exposure and outcome, coloc implements a Bayesian framework to derive the posterior probabilities of five hypotheses at a given locus: H_0_: There is no genetic association with either the exposure or the outcome; H_1_: There is genetic association with the exposure, but not the outcome; H_2_: There is genetic association with the outcome, but not the exposure; H_3_: There are distinct genetic associations with the exposure and with the outcome; and H_4_: There are shared genetic associations with both the exposure and the outcome.

The posterior probability of hypothesis H_4_ given the observed test statistics from GWAS is considered the colocalization probability. Importantly, coloc posterior probabilities are derived under the assumption that either (1) there exists at most one single causal variant shared between the exposure and the outcome, or (2) two distinct causal variants exist, one for the exposure and one for the outcome, within each locus[12]. This assumption may not be realistic for traits with a complex genetic architecture. Subsequently, extensions of coloc have been proposed to relax this assumption, such as an integration of the Sum of Single Effects (SuSiE) regression[13,14], referred to as coloc+SuSiE[15], and the pairwise conditional analysis and colocalization (PWCoCo) framework[16]. We recently developed another colocalization method, called SharePro[17], which uses a sparse projection shared between the exposure and the outcome to identify effect groups containing correlated variants, and assesses colocalization at the effect group level[17–19]. Several recent studies have leveraged pQTL-based MR analyses, combined with colocalization evidence generated using different methods, to identify candidate biomarkers or drug targets for complex diseases[16,20–28].

In previous simulation studies[17], all of these methods demonstrated appropriate false positive control, while SharePro achieved the highest accuracy in identifying shared causal variants, particularly when a locus contained multiple causal variants. However, a comprehensive benchmarking of the performance of these colocalization methods in real data analyses is still lacking. Therefore, in this study, we systematically evaluated the performance of coloc[12], coloc+SuSiE[15], PWCoCo[16], and SharePro[17] in providing colocalization evidence supporting MR-identified associations between circulating protein levels and key cardiometabolic traits, including diastolic blood pressure (DBP), systolic blood pressure (SBP), serum hemoglobin A1c (HbA1c), serum low-density lipoprotein cholesterol (LDL), and serum triglycerides (Tg). We investigated the performance of these methods, as well as their robustness to varying prior colocalization probabilities. Additionally, we examined whether colocalization evidence-supported associations agreed with results from exome-wide association studies (ExWAS) or implicated known drug targets for cardiometabolic diseases. This evaluation aims to provide insights into the reliability of these colocalization methods in the context of MR-guided target discovery, offering new lines of inquiry for precision medicine.

## Results

### 1. Colocalization evidence for protein-trait associations

An overview of the study design is presented in **Figure 1**. Cis-pQTL lead variants were selected as genetic instruments for the circulating levels of 1,535 proteins profiled in the Fenland study (**Supplementary Table S1**), which were individually tested for association with DBP, SBP, HbA1c, LDL, and Tg, respectively (**Supplementary Figures S1-S5**). The test statistics of MR analyses are available in **Supplementary Table S2**.

**Figure 1.**
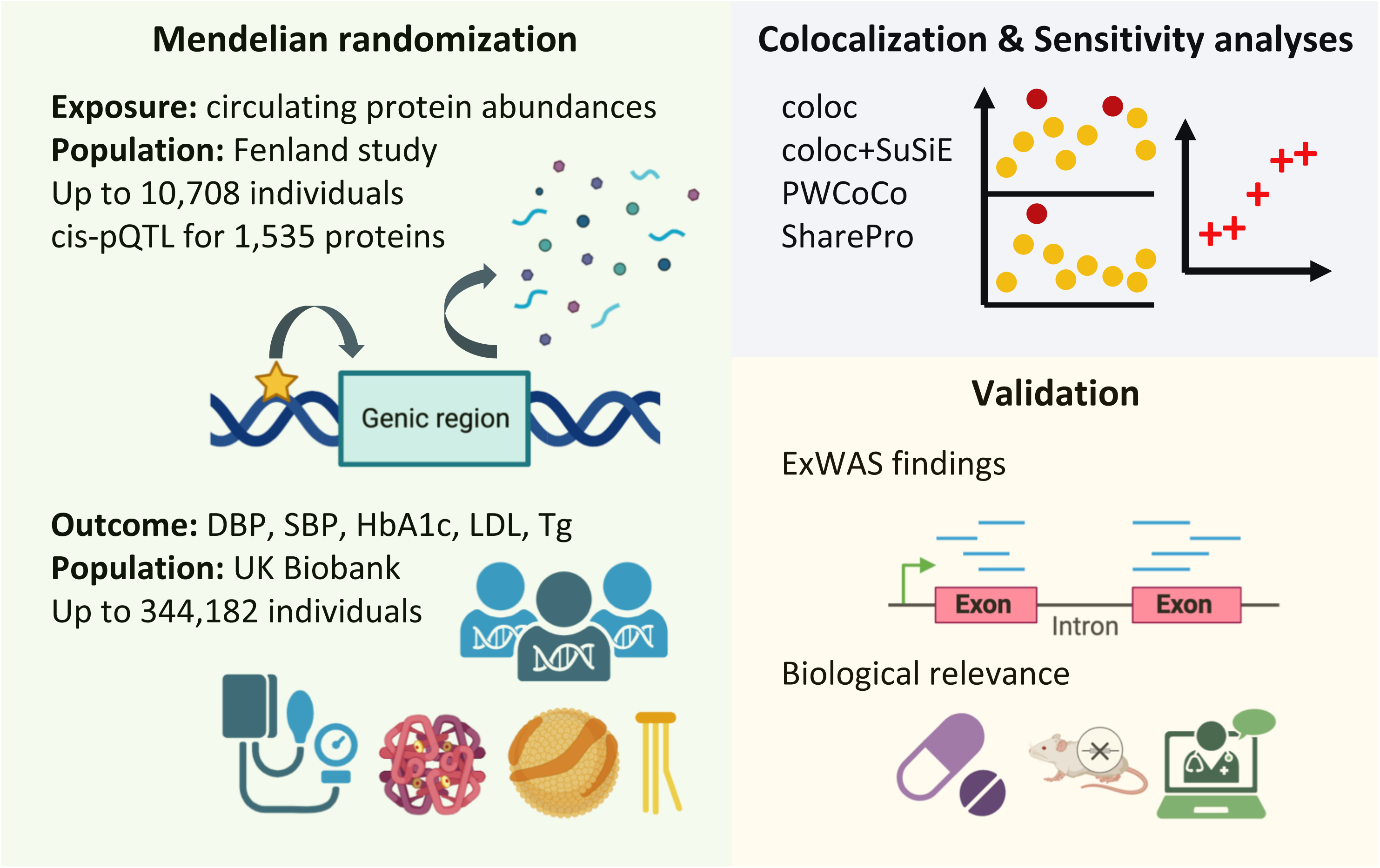
Study overview. Mendelian randomization was performed to assess the associations between circulating levels of 1,535 proteins and five cardiometabolic traits, including DBP, SBP, HbA1c, LDL, and Tg. Colocalization analyses and prior sensitivity analyses were performed using coloc, coloc+SuSiE, PWCoCo, and SharePro, respectively. Associations supported by colocalization evidence were examined with reference to ExWAS findings, successful drug targets, phenotypic changes in mouse mutants, and evidence in previous biomedical research. DBP: diastolic blood pressure; SBP: systolic blood pressure; HbA1c: hemoglobin A1c; LDL: low-density lipoprotein cholesterol; Tg: triglycerides; ExWAS: exome-wide association study.

Of the 7,675 tested associations, 611 non-MHC protein-trait pairs had a nominal p-value > 0.9. Colocalization analyses were first performed for these pairs with lack of evidence for associations using default settings of coloc, coloc+SuSiE, PWCoCo, and SharePro, respectively. As expected, none of the four methods supported the majority of most of these associations with strong or suggestive colocalization evidence. The only exception was that strong evidence of colocalization was found for the UBASH3B-HbA1c pair by SharePro (**Supplementary Table S3**). Interestingly, the selected instruments for the circulating level of UBASH3B were two variants in low LD, which exhibited opposing directions of effect in the MR analyses for HbA1c (**Supplementary Table S1** and **Supplementary Figure S6**). The effect groups separately tagged by these two genetic variants both appeared to be shared between the exposure and the outcome (**Supplementary Figure S6**), despite the inverse variance weighted estimate not demonstrating statistical significance. This inconsistency might be due to tissue-specific effects or measurement artifacts.

A total of 589 non-MHC protein-trait associations had an FDR < 0.05. This FDR threshold was close to a nominal p-value threshold of 4.1x10^-3^ (**Supplementary Table S2**). The minimum F-statistic of these 589 associations was 39.35 (**Supplementary Table S2**), suggesting a low risk of weak instrument bias. Furthermore, 201 associations were significant after a Bonferroni correction (p-value < 6.5x10^-6^). For proteins with at least three genetic instruments, MR estimates using alternative methods in sensitivity analyses were consistent with the primary MR estimates (**Methods** and **Supplementary Table S2**). No evidence of directional pleiotropy was detected (**Supplementary Table S2**).

Colocalization analyses were then performed for the 589 associations with an FDR < 0.05 using default settings (**Supplementary Table S4**). Under the single causal variant assumption, 109 (18.5%) associations were supported by strong colocalization evidence using coloc, including 81 (40.3%) of the 201 Bonferroni-significant associations (**Table 1**). coloc+SuSiE and PWCoCo moderately increased the number of associations supported by strong colocalization evidence to 127 (21.6%) and 126 (21.4%), including 94 (46.8%) and 92 (45.8%) of the Bonferroni-significant associations, respectively (**Table 1**). Colocalization probabilities inferred using coloc, coloc+SuSiE, and PWCoCo were largely consistent, apart from increments as a result of modelling multiple causal effects (**Supplementary Figures S7-S11**). In contrast, there were differences between colocalization probabilities inferred using SharePro and the other three methods (**Supplementary Figures S7-S11**). Overall, SharePro identified strong colocalization evidence for 179 (30.4%) protein-trait associations (**Table 1**). Importantly, 160 of these associations withstood Bonferroni correction, constituting 79.6% of all Bonferroni-significant associations (**Table 1**). Notably, only 27 (13.4%) Bonferroni-significant associations were not supported by strong or suggestive colocalization evidence using SharePro, compared to 108 (53.7%), 94 (46.8%), and 100 (49.8%) using coloc, coloc+SuSiE, and PWCoCo, respectively (**Table 1**). Consistent across all five traits, although SharePro supported the fewest associations with an FDR < 0.05 that did not reach the Bonferroni-corrected significance threshold, it supported the most Bonferroni-significant associations (**Table 1**).

**Table 1.**
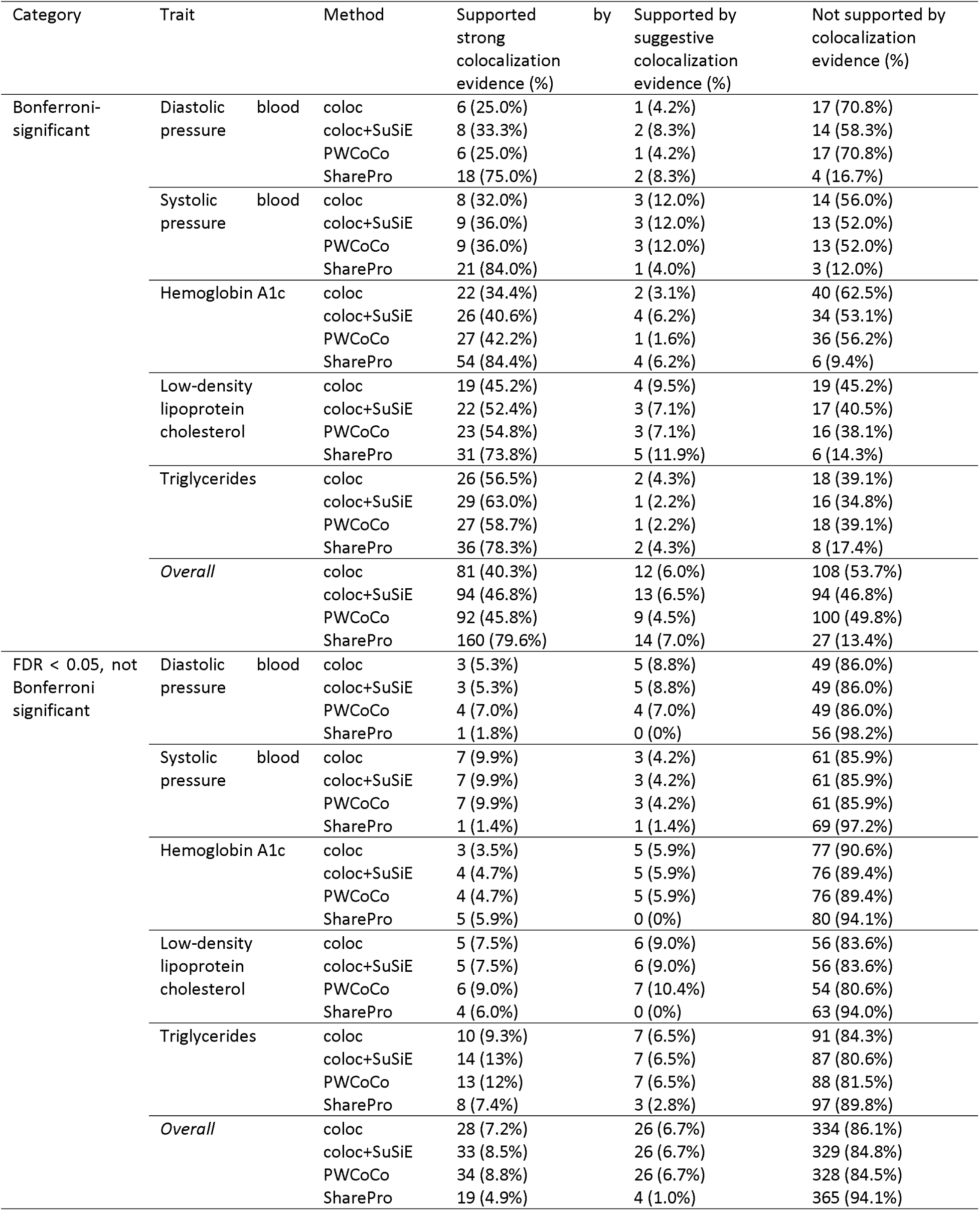
Numbers of protein-cardiometabolic trait associations supported by colocalization evidence.

All four methods were designed to be computationally efficient[12,15–17]. Among them, coloc and SharePro were faster than coloc+SuSiE and PWCoCo, while PWCoCo was the most memory-efficient (**Supplementary Table S5**).

### 2. Prior sensitivity analyses

Since the prior colocalization probability can be important in colocalization analyses, sensitivity analyses with different values of the priors were performed (**Methods**). While the impact of the prior colocalization probability was as anticipated, it is noteworthy that as the prior increased from 1.0x10^-7^ to 1.0x10^-3^, the number of protein-trait associations supported by SharePro was less variable than those suppoted by coloc, coloc+SuSiE, and PWCoCo (**Figure 2A**). Importantly, although some associations supported by coloc, coloc+SuSiE, or PWCoCo with the default prior of 1.0x10^-5^ were not simultaneously supported by SharePro with its default prior, all such associations were at least supported by suggestive colocalization evidence using SharePro when the prior increased to 1.0x10^-3^ (**Figure 2B-D**). In addition, SharePro was able to support more of these associations when the colocalization probabilities decreased dramatically using the other methods with the weakest prior of 1.0x10^-7^ (**Figure 2B-D**). In contrast, more than 25% of the associations supported by SharePro with the default prior had a colocalization probability < 0.5 using coloc, coloc+SuSiE, or PWCoCo with the strongest prior of 1.0x10^-3^ (**Figure 2E**). This trend was consistent across all five traits (**Supplementary Figures S12-S16**). We identified 33 protein-trait associations that were exclusively supported by SharePro, whereas no association was exclusively supported by coloc, coloc+SuSiE, or SharePro (**Methods** and **Supplementary Table S4**).

**Figure 2.**
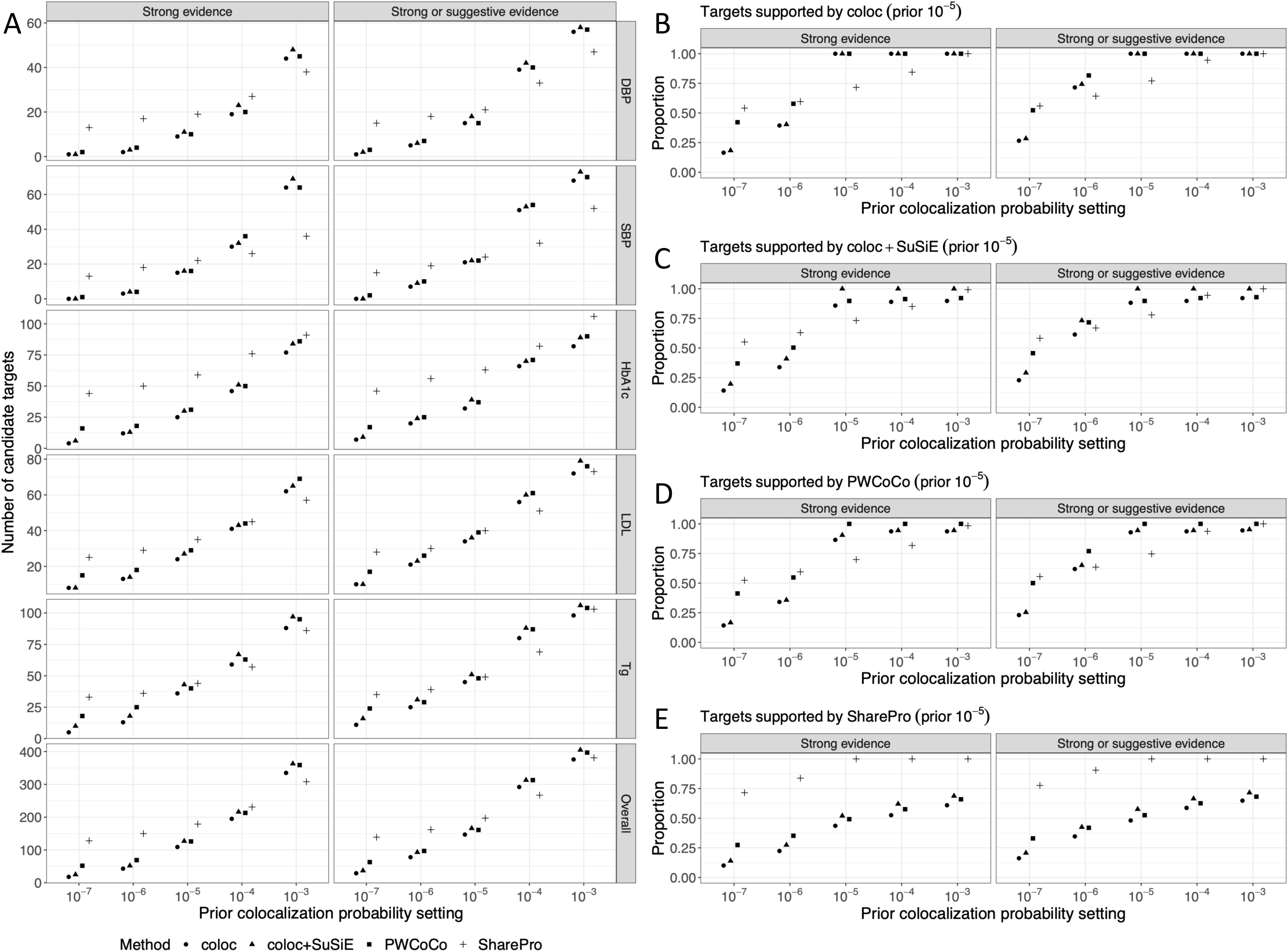
Results of prior sensitivity analyses. (A) Protein-cardiometabolic trait associations supported by colocalization evidence with different prior settings. The number of candidate targets supported by SharePro was the least variable across different prior colocalization probabilities. For protein-trait associations supported by strong colocalization evidence using (B) coloc, (C) coloc+SuSiE, (D) PWCoCo, and (E) SharePro with the default prior of 1.0x10-5, respectively, the proportions of these associations supported by each method using different prior settings are summarized. The observed trend is consistent across all five traits, as depicted in Supplementary Figures S12-S16.

### 3. Agreement between colocalization evidence and rare variant-based gene-level associations

We next examined whether findings in MR and colocalization analyses agreed with significant gene-level associations detected in ExWAS for the five traits (**Methods**). Fourteen gene-trait associations involving HbA1c, LDL, and Tg demonstrated suggestive evidence of association in ExWAS with a p-value < 1.0x10^-5^, including nine genome-wide significant associations with a p-value < 5.0x10^-8^ (**Supplementary Table S6**). The most significant associations implicated genes with well-characterized roles in the metabolism of LDL and Tg, such as *APOC3*[29–31], *APOB*[32–34], and *PCSK9*[32,35–37] (**Supplementary Table S6**). For each of these 14 associations, the MR-estimated effect of the genetically decreased circulating protein level on the respective cardiometabolic trait had the same direction as the collapsing model containing rare variants predicted to be deleterious (**Figure 3**).

**Figure 3.**
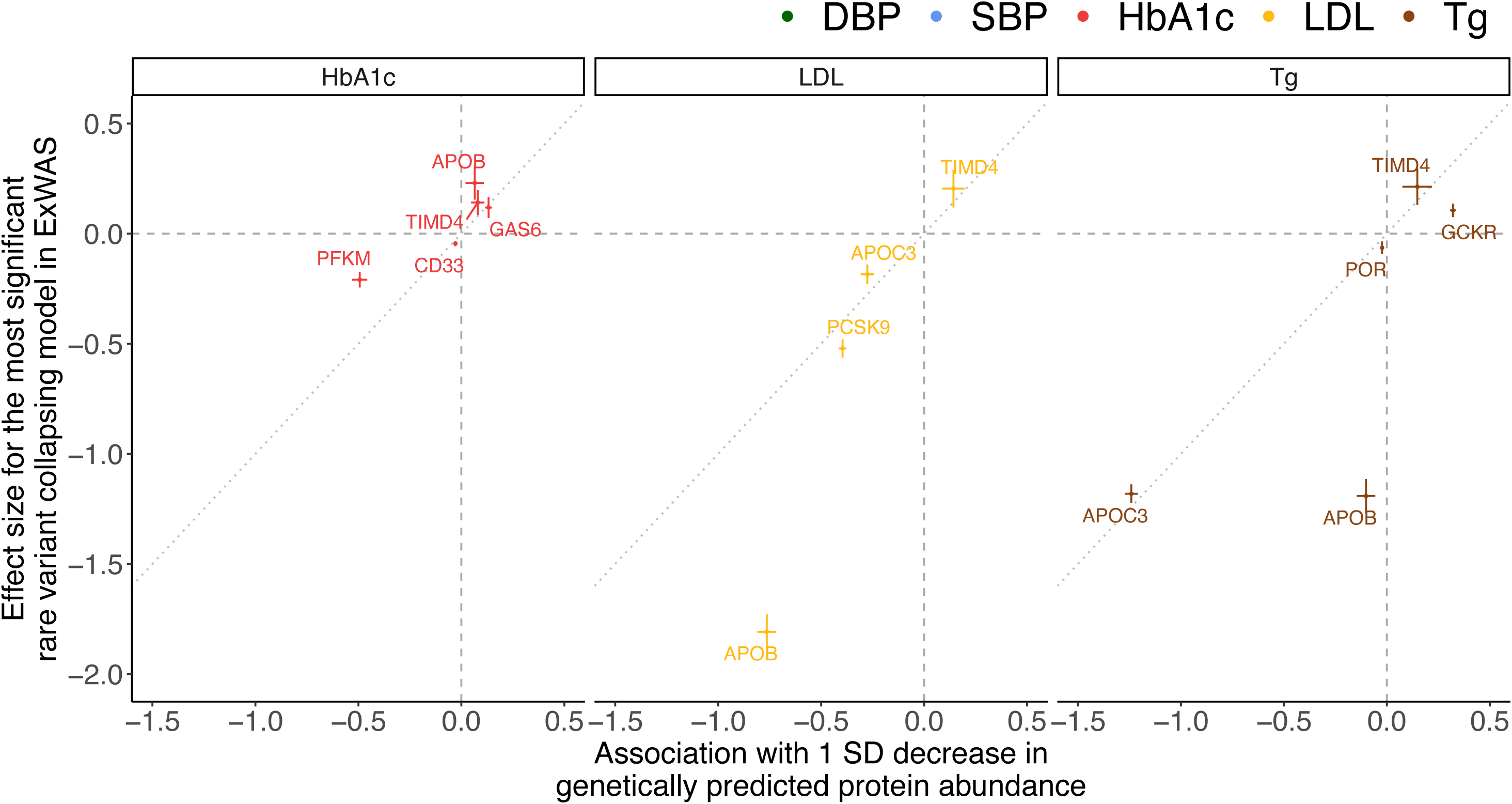
Comparison of estimated effects in rare variant collapsing analyses and Mendelian randomization. The estimated average difference in the outcome between individuals with the variants and those without for the most significant rare variant collapsing model is compared with the estimated change in outcome associated with a one SD decrease in genetically predicted circulating protein level. The dotted lines have a slope of 1. Genes with a p-value < 1.0x10^-5^ in ExWASs are illustrated. SD: standard deviation; ExWAS: exome-wide association study.

Using default settings, SharePro supported 13 of the 14 associations with strong colocalization evidence, compared to 10 by coloc, 11 by coloc+SuSiE, and 11 by PWCoCo (**Figure 4**). Despite coloc showing a low colocalization probability, coloc+SuSiE, PWCoCo, and SharePro supported the PFKM-HbA1c association (**Figure 4**), wherein PFKM is a muscle phosphofructokinase and regulatory enzyme in glycolysis[38,39]. SharePro supported the APOB-Tg and POR-Tg associations with the default prior (**Figure 4**), although the other methods also supported these associations with a stronger prior (**Supplementary Figure S17**). Colocalization analyses for the APOB-HbA1c association were sensitive to the selected prior, and no method provided colocalization evidence with the default prior (**Figure 4** and **Supplementary Figure S17**).

**Figure 4.**
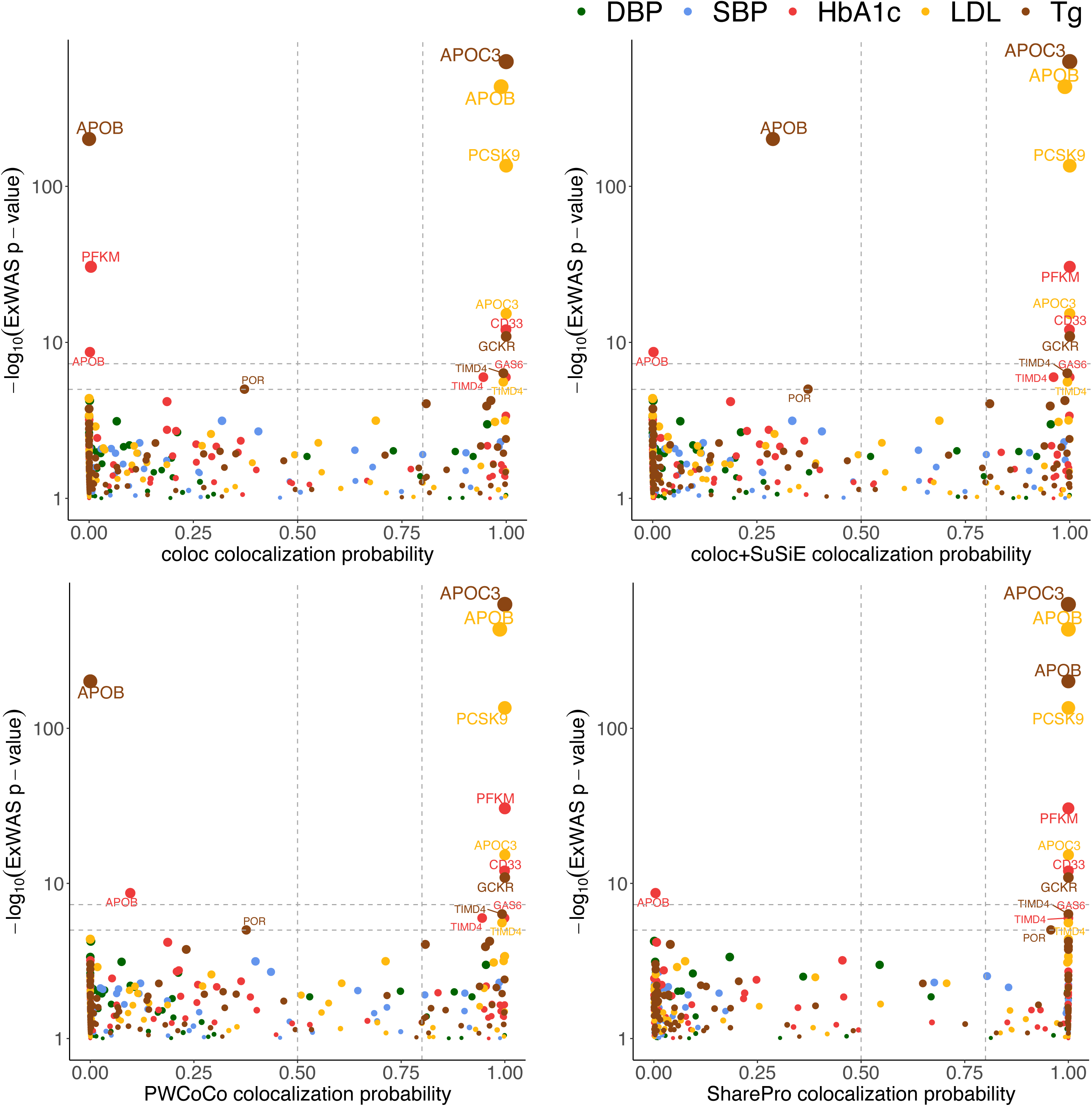
Agreement between colocalization evidence and ExWAS findings. The significance of each gene in the ExWAS of each trait is plotted against the inferred colocalization probability of the corresponding protein-trait association for each colocalization method. Two horizontal dashed lines represent p-value thresholds of 1.0x10^-5^ and 5.0x10^-8^, respectively. Two vertical dashed lines represent colocalization probability thresholds of 0.5 and 0.8, respectively. Genes with a p-value < 0.1 in ExWASs are illustrated. Sensitivity analyses are illustrated in Supplementary Figure S17. ExWAS: exome-wide association study.

### 4. Colocalization evidence for known drug targets

We then assessed colocalization evidence for known drug targets in the DrugBank database (**Methods** and **Supplementary Table S7**). Among the 22 known drug targets under investigation for at least one of the five cardiometabolic traits, coloc and PWCoCo with the default prior identified strong colocalization evidence supporting the associations between the circulating levels of five proteins and the relevant cardiometabolic traits (**Methods** and **Figure 5**), namely PCSK9 (hypercholesterolemia, myocardial infarction and stroke), GNMT (myocardial infarction and stroke), GSS (myocardial infarction and stroke), PLAU (heart failure), and SULT2A1 (atrial fibrillation). coloc+SuSiE further supported the prothrombin (F2)-Tg association, which implicates drug targets for myocardial infarction and stroke, with strong colocalization evidence (**Figure 5** and **Supplementary Table S7**). SharePro was able to identify strong colocalization evidence for all of these six drug targets as well as three additional targets (**Figure 5**), including ALDH2 (hypertension and heart failure), NPPB (hypertension and heart failure), and PCSK1 (diabetes). Genetic associations with the circulating levels of ALDH2, NPPB, PCSK1, and F2 and the corresponding traits are depicted in **Figure 6**. While multiple causal variants likely exist within the same loci, variants in high LD (r^2^ > 0.8) with the genetic instruments of circulating protein levels demonstrated genome-wide significance in the GWAS of the respective trait (**Figure 6**). Notably, the ALDH2-DBP, NPPB-DBP, and NPPB-SBP associations were exclusively supported by SharePro as the inferred colocalization probabilities using other methods were below 0.5 with the strongest prior (**Supplementary Figure S18**).

**Figure 5.**
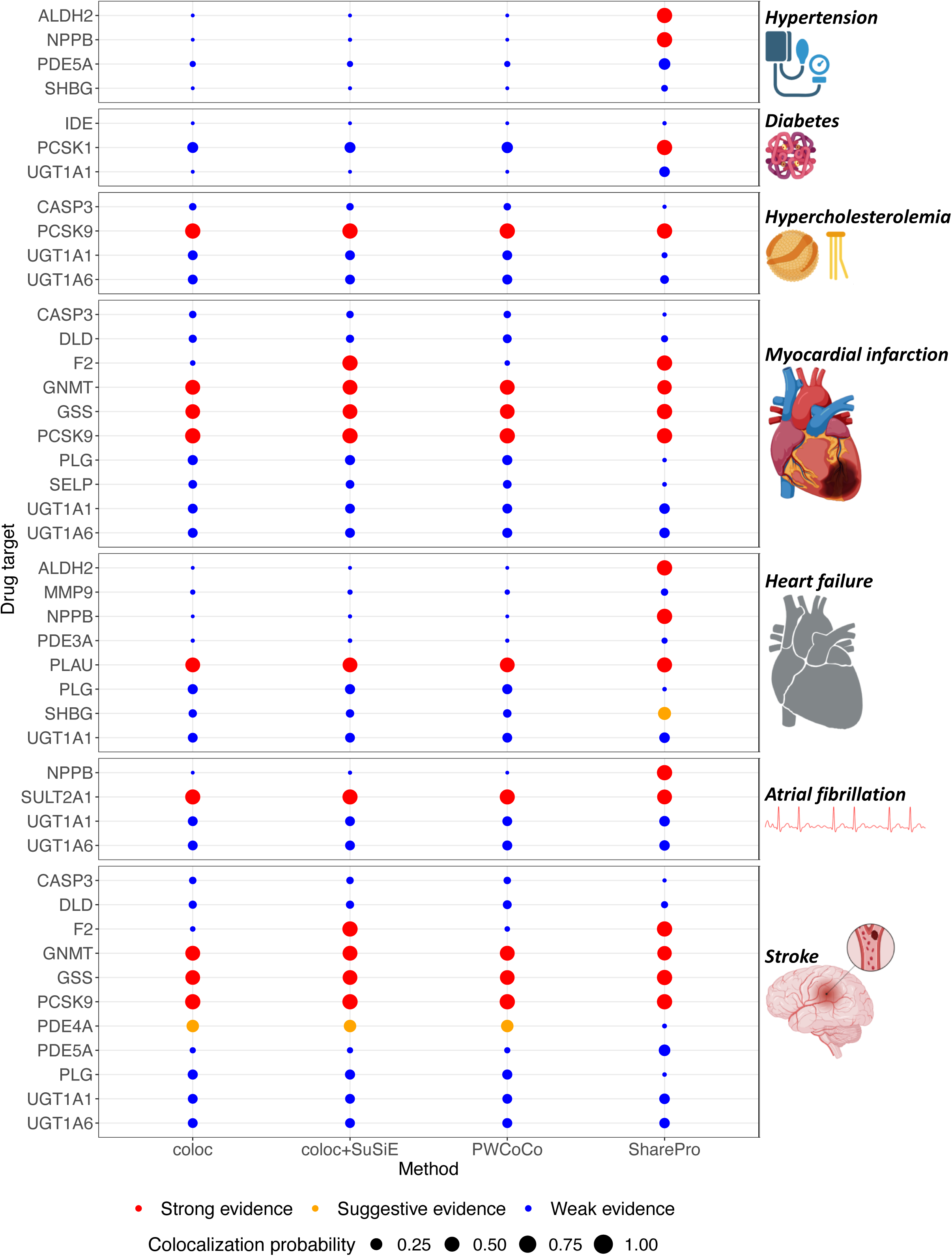
Associations supported by colocalization evidence implicate successful drug targets. For each method, colocalization evidence is summarized for each drug target of the seven cardiometabolic diseases, based on the maximal colocalization probability across all associations between circulating protein levels and relevant cardiometabolic traits, as detailed in Methods.

**Figure 6.**
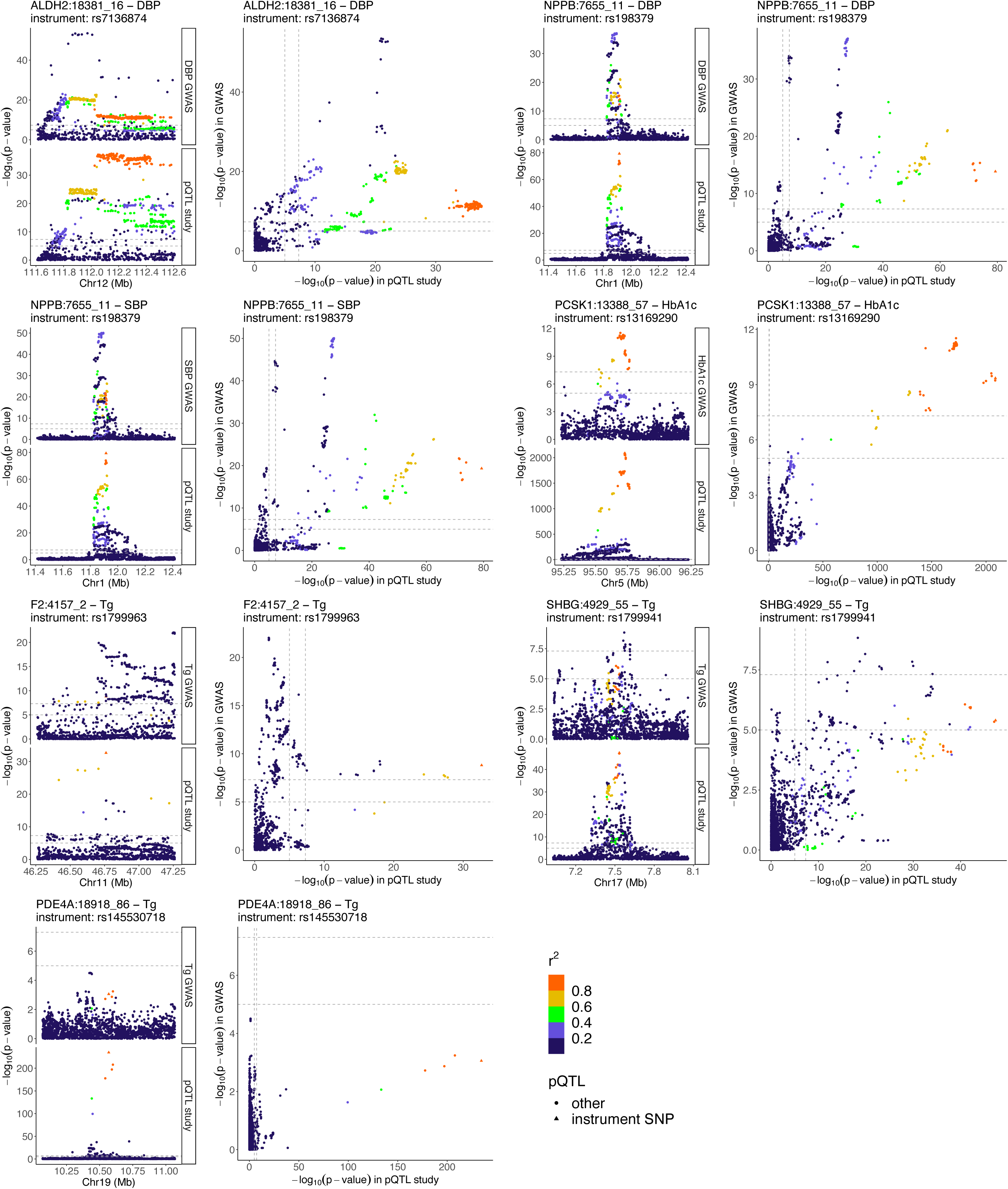
Illustration of genetic associations involving ALDH2, NPPB, PCSK1, F2, SHBG, and PDE4A with relevant traits. The lead variants of cis-pQTLs used as genetic instruments are indicated. Genetic variants located in a ±500kb window centered around each genetic instrument are plotted with their significance in respective studies, and colored by the magnitude of correlation (linkage disequilibrium r^2^) with the corresponding instrument. Two horizontal dashed lines and two vertical dashed lines represent p-value thresholds of 1.0x10^-5^ and 5.0x10^-8^, respectively. Protein names include SOMAmer identifiers. Sensitivity analyses are illustrated in Supplementary Figure S18. Cis-pQTL: cis-protein quantitative trait locus.

Furthermore, with the default prior, SharePro identified suggestive colocalization evidence for an association between SHBG and Tg, implicating a drug target for heart failure (**Figure 5** and **Supplementary Table S7**). On the other hand, suggestive colocalization evidence for the PDE4A-Tg association, which implicates a drug target for stroke, were identified by coloc, coloc+SuSiE, and PWCoCo, but not by SharePro (**Figure 5** and **Supplementary Table S7**). However, these results were sensitive to the prior settings (**Supplementary Figure S18**).

### 5. Protein-trait associations exclusively supported by SharePro

Finally, we investigated the plausibility of the 33 protein-trait associations exclusively supported by SharePro (**Methods** and **Supplementary Figure S19**). Of these, 20 associations involved DBP, SBP, LDL, or Tg, and their colocalization evidence was reassessed using SharePro based on larger, meta-analyses of European ancestry-specific GWAS. As a result, 19 associations remained supported by strong colocalization evidence, and one association was supported by suggestive colocalization evidence (**Supplementary Table S8**). Systematic evaluation of the risk of horizontal pleiotropy was conducted, leading to the exclusion of 25 associations that could be subject to a high risk of horizontal pleiotropy (**Methods** and **Supplementary Tables S9-S11**). The risk of horizontal pleiotropy was mild for the remaining eight associations (**Figure 7**).

**Figure 7.**
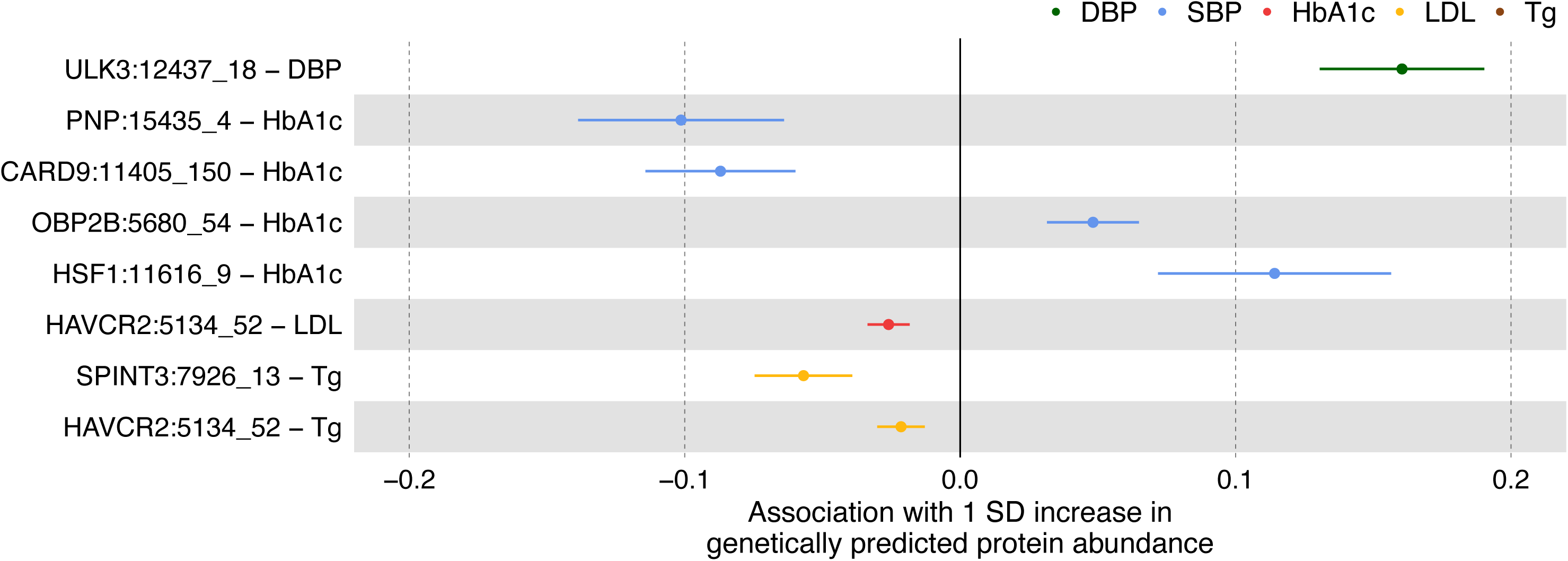
Protein-trait associations exclusively supported by SharePro. Eight protein-cardiometabolic trait associations identified through Mendelian randomization that were not subject to a high risk of horizontal pleiotropy. The estimated effects of a one SD increase in genetically predicted circulating protein levels on the standardized outcomes are illustrated. SD: standard deviation.

These genetic associations are illustrated in **Figure 8**. Again, variants in high LD (r^2^ > 0.8) with the genetic instruments of circulating protein levels also demonstrated genome-wide significance in the GWAS of the corresponding trait for all of these protein-trait associations (**Figure 8**), although there could be other non-colocalizing causal variants underlying the trait within the same loci. Phenotypic changes in mouse mutants of homologous genes are summarized in **Supplementary Table S12**.

**Figure 8.**
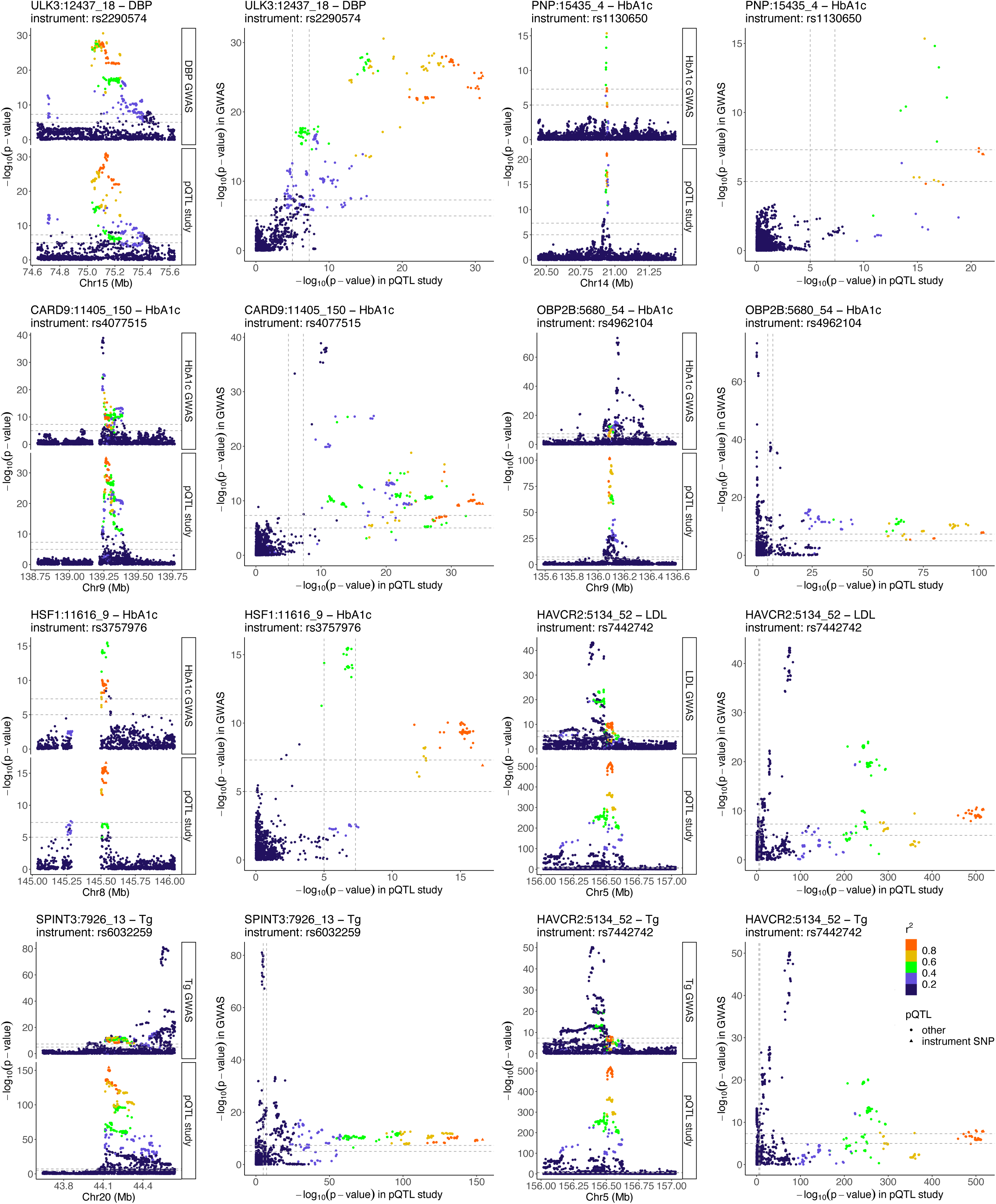
Illustration of genetic associations underlying protein-trait associations exclusively supported by SharePro. The lead variants of cis-pQTLs used as genetic instruments are indicated. Genetic variants located in a ±500kb window centered around each genetic instrument are plotted with their significance in respective studies, and colored by the magnitude of correlation (linkage disequilibrium r^2^) with the corresponding instrument. Two horizontal dashed lines and two vertical dashed lines represent p-value thresholds of 1.0x10^-5^ and 5.0x10^-8^, respectively. Protein names include SOMAmer identifiers. Sensitivity analyses are illustrated in Supplementary Figure S19. Cis-pQTL: cis-protein quantitative trait locus.

## Discussion

In this study, we benchmarked the performance of four Bayesian colocalization methods, including coloc, coloc+SuSiE, PWCoCo, and SharePro, in the context of supporting circulating protein-cardiometabolic trait associations identifying through MR.

We demonstrated that SharePro supported substantially more (79.6%) Bonferroni-significant associations with strong colocalization evidence than coloc (40.3%), coloc+SuSiE (46.8%), and PWCoCo (45.8%). On the other hand, coloc, coloc+SuSiE, and PWCoCo supported more associations with an FDR < 0.05 that were not Bonferroni-significant. Furthermore, MR-identified protein-trait associations that were supported by SharePro were more likely to agree with gene-level associations detected in ExWAS and implicated more known drug targets for cardiometabolic diseases compared to the other methods. Notably, only SharePro supported the associations between Tg and APOB and between blood pressure and NPPB. The *APOB* gene was significantly associated with Tg in ExWAS and is clinically actionable for familial hypercholesterolemia[32–34]. NPPB, known as B-type natriuretic peptide, is a well-characterized regulator of vascular resistance and blood pressure and a target of anti-hypertensive drugs[40–42].

By design, there are notable differences between these colocalization methods. Both coloc+SuSiE and PWCoCo employ a two-step strategy for colocalization assessment. Specifically, coloc+SuSiE first performs statistical fine-mapping to identify 95% credible sets using SuSiE (a credible set is the smallest set of variants such that there is a 95% probability that the true causal variant is within this set)[13–15], while PWCoCo first performs conditional and joint analyses to identify conditional independent genetic associations[16]. This first step is conducted separately for the exposure and the outcome. Then, colocalization analyses are performed for each pair of credible sets in coloc+SuSiE and each pair of conditional independent signals in PWCoCo. Hence, for both methods, colocalization assessment in the second step might be biased by the uncertainty in the first step. In contrast, SharePro integrates LD modelling and colocalization assessment through a shared sparse projection between genetic associations with the exposure and the outcome, mitigating the aforementioned bias and possibly better aligning genetic signals[17]. Importantly, SharePro requires that both the exposure GWAS and the outcome GWAS should be sufficiently powered (i.e., with a large sample size) in order to recognize effect groups and assess colocalization evidence at the effect group level[17]. As a result, MR-identified associations that had an FDR < 0.05 but were not Bonferroni-significant were less likely to be supported by SharePro, since most of the genetic associations with the cardiometabolic traits did not reach genome-wide significance within the respective loci.

Prior sensitivity analyses are important for assessing the robustness of colocalization analyses[43]. In this study, SharePro demonstrated the strongest robustness to the choice of prior colocalization probability. Remarkably, associations supported by strong evidence using coloc, coloc+SuSiE, or PWCoCo with the default prior were more likely to be supported using SharePro when the prior was weakened. Conversely, SharePro could support all of these associations when the prior was increased. It is important to note that using a high prior carries a risk of false positives[17]. Therefore, interpretation of results under a high prior should be made with caution. The sensitivity analyses are intended to evaluate the robustness of findings across a range of prior settings rather than to endorse the use of elevated priors in practice.

We identified eight protein-cardiometabolic trait associations exclusively supported by SharePro that were not subject to a high risk of horizontal pleiotropy. Several protein-coding genes showed biological relevance in mouse mutants. In particular, increased circulating levels of HSF1 were predicted to increase HbA1c, aligning with the observation that knockout mice of the heat shock transcription factor 1 (*Hsf1*) gene have decreased circulating glucose levels compared to wild types[44]. The regulatory role of HSF1 in glucose homeostasis has also been suggested in studies of human cell lines and mice[45–49]. Notably, HSF1 likely functions as an intracellular protein and may be released into circulation as a result of cell death. Therefore, it is likely more promising to explore circulating HSF1 as a biomarker rather than a direct drug target. Furthermore, increased circulating levels of HAVCR2, also known as T-cell immunoglobulin and mucin-domain containing-3, were predicted to decrease LDL and Tg. HAVCR2 is an important regulator of T-cell responses[50], and is preferentially expressed on Th1 helper T-lymphocytes that are known to promote atherogenesis[51,52], although it is also expressed on other T-cell subsets whose roles in the development of atherosclerosis are less well defined[51,52]. Interestingly, blockade of HAVCR2 in low-density lipoprotein receptor-deficient mice could exacerbate atherosclerosis[53], and monocyte expression of *HAVCR2* has been found to be lower in patients with coronary artery disease than in healthy control subjects[54]. These MR-identified associations warrant further investigations.

However, the interpretation of these distinctive findings still requires extra caution. The genetic instruments used for the circulating levels of these proteins all demonstrated a mild risk of horizontal pleiotropy due to their associations with the expression level or alternative splicing of neighboring genes. For example, the genetic instrument of the circulating level of ULK3 (rs2290574) has been associated with the gene expression level of a nearby gene *CSK*. In a small rodent study, changes in blood pressure have been observed in mice treated with an siRNA directed against *Csk*, but not in *Ulk3* siRNA-treated mice, although the sample size was small[55]. The increased risk of horizontal pleiotropy is expected for these associations exclusively supported by one colocalization method, given the evident existence of multiple causal variants and potentially increased LD complexity compared to other loci. Therefore, further evidence, particularly from functional validation studies, is necessary to confirm the biological plausibility of such targets in future studies.

Taken together, we suggest that in practice, multiple methods should be applied to assess colocalization evidence, accompanied by prior sensitivity analyses. When results obtained using different methods show discrepancies, interpretation of MR-identified associations should consider various factors, including model assumptions of different colocalization methods, distinct LD structures, statistical power of GWAS, as well as biological plausibility.

Important limitations should be noted in this study. First, there is no genome-wide gold standard dataset containing unequivocal true positive and true negative protein-trait associations. This lack of definitive ground truth makes it challenging to precisely benchmark the sensitivity and false positive rates of the colocalization methods evaluated. Although we incorporated other types of evidence, these approaches cannot fully capture true causal relationships. Method performance may also be affected by unmeasured confounding, incomplete coverage of the proteome and phenome, and inherent limitations of the datasets used. Furthermore, our benchmarking focused specifically on circulating proteins in relation to five cardiometabolic traits. While this focus was intentional, given the strong biological relevance of cardiometabolic diseases to circulating protein levels, it may limit the generalizability of our findings to other types of molecular traits or complex phenotypes. In fact, extending this framework to traits or diseases less directly linked to circulating proteins would introduce additional complexity and require tailored methodological considerations. Future studies expanding to a broader range of traits and diseases, along with experimental validation and well-curated reference datasets, will be critical for fully evaluating and maximizing the potential colocalization methods. Second, our analyses have not encompassed all statistical methods available for colocalization analyses, such as eCAVIAR[56] and fastENLOC[57,58]. eCAVIAR performs statistical fine-mapping within a locus separately for the exposure and the outcome[56]. However, unlike coloc+SuSiE and SharePro, fine-mapping in eCAVIAR necessitates a computationally expensive search through possible causal configurations[56]. Colocalization probabilities are then calculated at the variant level rather than the credible set or locus level. coloc+SuSiE, PWCoCo, and SharePro have been shown to achieve a higher accuracy than eCAVIAR in previous studies[13,17]. fastENLOC adopts a data-driven approach to estimate the enrichment of genetic associations in QTLs[57,58]. These enrichment parameters are subsequently used to specify a fixed prior colocalization probability for all loci under investigation[57,58]. While this approach may be effective for understanding the overall genetic architecture of complex traits, we posit that locus-specific colocalization evidence combined with agnostic prior sensitivity analyses could better identify colocalized associations that are detected in GWAS. Furthermore, our primary analyses were restricted to populations of European ancestry based in the United Kingdom, since all colocalization methods that model LD structures require an LD reference panel that is similar to the GWAS populations. Potential LD mismatch between the LD reference panel and GWAS populations, even within the same ancestry, can substantially reduce the accuracy of existing colocalization methods[59]. Consequently, we did not fully utilize the largest GWAS meta-analyses of cardiometabolic biomarkers and disease outcomes[60–64], which have included heterogeneous populations that can possess unique LD structures. Advocacy for releasing in-sample LD information from large GWAS and developing novel algorithms with enhanced tolerance to LD mismatch may further improve colocalization analyses in diverse populations in the future. Finally, since sex differences are known to affect the risk and progression of cardiometabolic diseases[65], future work incorporating sex-stratified analyses or interaction models[18] may provide additional insights into potentially sex-specific mechanisms and improve the precision of target discovery efforts.

In summary, our findings suggest that SharePro is an accurate and robust Bayesian colocalization method for supporting statistically significant associations identified through pQTL-based MR. By refining colocalization evidence through the consideration of results obtained using multiple methods and from sensitivity analyses, the yield of biomarker and drug target discovery programs may be substantially increased for cardiometabolic conditions and other complex diseases.

## Materials and Methods

### 1. Genome-wide association studies

We obtained summary statistics from GWAS of circulating protein levels and the five cardiometabolic traits. Circulating proteins in plasma were quantified in the Fenland study using the SomaLogic SomaScan v4 assay[6] that measured a total of 4,979 SOMAmer protein targets. This study comprised 10,708 unrelated individuals of European ancestry, aged 29-64 years at recruitment, from Cambridgeshire, United Kingdom. The cohort and the proteo-genomic study have been detailed previously[6,66]. Genome-wide association analyses of the inverse normal transformed circulating level of each protein were conducted. Distinct lead variants associated with the circulating protein levels were identified through conditional analyses, with a Bonferroni-corrected genome-wide significance threshold of p-value < 1.0x10^-11^. Distinct variants located within 500kb of the gene body of each protein-coding gene were considered cis-pQTL lead variants.

The UK Biobank recruited approximately 500,000 individuals aged 40-69 years at multiple recruitment centers in the United Kingdom[67]. Various physical measures and biological samples were collected during the baseline assessment. Genome-wide association analyses of the inverse normal transformed DBP (N = 340,162), SBP (N = 340,159), HbA1c (N = 344,182), LDL (N = 343,621), and Tg (N = 343,992) were conducted by the Neale Lab (https://github.com/Nealelab/UK_Biobank_GWAS)[68]. The genetic ancestry of each participants was derived based on reference data from the 1000 Genomes Project[69] through the clustering of the leading genetic principal components[67]. In this study, we focused on the European ancestry-specific GWAS results to minimize LD mismatch. There was no known sample overlap between the Fenland study and the UK Biobank.

### 2. Mendelian randomization and sensitivity analyses

We performed two-sample MR to assess the association between the circulating level of each protein and each of the five cardiometabolic traits. We selected cis-pQTL lead variants detected in the Fenland study as genetic instruments. We did not utilize trans-pQTL variants due to an increased risk of horizontal pleiotropy involving other genes and pathways. If a cis-pQTL lead variant was not present in the cardiometabolic trait GWAS, a proxy in high LD with this variant was used as the genetic instrument when available (squared Pearson correlation coefficient, r^2^ > 0.8; an r^2^ = 1 indicates perfect correlation between two variants, while an r^2^ = 0 indicates no correlation). Proxy searches were performed using the LDlink R package[70] based on reference data from the 1000 Genomes Project non-Finnish European ancestry populations. Summary statistics were harmonized using the default settings of the TwoSampleMR R package version 0.5.6[71], aligning effect alleles between the exposure and outcome. Forward strand alleles were inferred based on allele frequency information. Palindromic variants with a minor allele frequency > 0.42 were excluded to avoid strand mismatches. In total, 1,535 proteins with at least one available genetic instrument were included in the following MR analyses.

Primary MR estimates were obtained using the Wald ratio method for proteins with one genetic instrument and the inverse variance weighted method for proteins with at least two genetic instruments. Associations with a Benjamini-Hochberg-adjusted p-value (false discovery rate, FDR) < 0.05 were considered significant, accounting for a total of 7,675 (1,535 proteins x 5 traits) tests. Furthermore, associations with a p-value < 0.05/7,675 = 6.5x10^-6^ were considered Bonferroni-significant.

For proteins with at least three genetic instruments, we performed sensitivity analyses using the weighted mode, weighted median, and penalized weighted median methods[72–74]. We assessed whether the estimated direction and magnitude of effect were consistent using these different methods. We tested for directional horizontal pleiotropy using the MR-Egger method[75], where a nominally significant MR-Egger intercept (p-value < 0.05) was considered evidence of directional pleiotropy.

For each test, we calculated the F-statistic, where an F-statistic < 10 is indicative of weak instrument bias[76]. MR analyses were performed using the TwoSampleMR R package version 0.5.6[71].

### 3. Colocalization and prior sensitivity analyses

We performed colocalization analyses for two groups of protein-trait pairs: (1) those with a nominal p-value > 0.90 in primary MR analyses; and (2) those with an FDR < 0.05 in primary MR analyses. We excluded proteins coded by genes that are located in the major histocompatibility complex (MHC) region (genome assembly GRCh37.p13), because of its high variability and complex LD structure[77]. Most of the unassociated protein-trait pairs in group (1) could serve as negative controls since it is unlikely for these genomic loci to harbor shared genetic associations with both the circulating protein levels and the traits. However, exceptions can arise when multiple genetic instruments have opposing effect directions, leading to the possibility of the inverse variance weighted estimates not demonstrating statistical significance. For each protein-trait pair, we gathered GWAS summary statistics of all variants that are within 500kb of the cis-genetic instruments. We randomly selected 30,000 unrelated European ancestry individuals from the UK Biobank as the LD reference panel.

Posterior colocalization probabilities were initially inferred using the default priors of coloc, coloc+SuSiE, PWCoCo, and SharePro, respectively. For coloc, coloc+SuSiE, and PWCoCo, the default priors are p_1_ (prior probability of the exposure having a causal variant) = 1.0x10^-4^, p (prior probability of the outcome having a causal variant) = 1.0x10^-4^, and p_12_ (prior probability of the exposure and the outcome sharing the same causal variant) = 1.0x10^-5^. Among these priors, p_1_ and p_2_ are well-justified, broadly corresponding to a 99% belief that a true causal variant exists when there is a genome-wide significant association (i.e. p-value < 5.0x10^-8^)[43], while p_12_ exerts a major influence on the posterior colocalization probability. SharePro uses a default prior colocalization probability of 1.0x10^-5^, which is analogous to setting p = 1.0x10^-5^.

For coloc+SuSiE and SharePro, we set the maximal number of causal effects in each locus (K) to be 10. Importantly, when coloc+SuSiE or SharePro identified two or more credible sets or effect groups (K ≥ 2), we required that each credible set or effect group must be tagged by at least one genetic instrument of the circulating protein level, defined as the lead variant of the credible set or effect group being in high LD (r^2^ > 0.8) with a genetic instrument. Credible sets or effect groups that were not tagged by any genetic instruments through LD were discarded. For PWCoCo, we set the maximal number of conditionally independent lead variants to be 10.

Following practical recommendations from previous studies[15,16], for coloc+SuSiE, the locus-wise posterior colocalization probability was obtained by taking the maximum of coloc-inferred posterior probability (assuming a single causal variant) and all coloc+SuSiE-inferred credible set-level posterior colocalization probabilities; for PWCoCo, the locus-wise posterior colocalization probability was obtained by taking the maximum of coloc-inferred posterior probability (without conditional analyses) and all conditional posterior colocalization probabilities. For SharePro, the locus-wise posterior colocalization probability was obtained by taking the maximum of posterior colocalization probability inferred under the single causal effect group assumption (K = 1, analogous to coloc) and all effect group-level posterior colocalization probabilities for K ≥ 2[17]. Runtime and peak memory usage were compared across all methods using a single 2 x Intel E5-2683 v4 Broadwell @ 2.1GHz CPU core on Digital Research Alliance of Canada infrastructure.

To evaluate the impact of varying prior colocalization probabilities[43], we repeated colocalization analyses by varying the prior colocalization probability in {1.0x10^-3^, 1.0x10^-4^, 1.0x10^-5^ (default), 1.0x10^-6^, 1.0x10^-7^}. Notably, setting strong priors may lead to false positives, as demonstrated in prior work[17]. In this study, higher prior colocalization probabilities were used solely to characterize how different methods behave under varying prior settings and to assess the robustness of colocalization evidence. These sensitivity analyses should not override previously noted caution regarding the potential for inflated colocalization probabilities.

In all colocalization analyses, a colocalization probability > 80% was considered strong evidence of colocalization, while a colocalization probability > 50% was considered suggestive evidence of colocalization[12,16]. A protein-trait association was exclusively supported by one method if this method could provide strong colocalization evidence with the weakest prior of 1.0x10^-7^, while none of the other methods could provide suggestive colocalization evidence with the strongest prior of 1.0x10^-3^.

### 4. Gene-level associations with selected cardiometabolic traits

We assessed whether MR and colocalization evidence agreed with findings of collapsing analyses of rare variants using whole-exome sequencing data from the UK Biobank, as the latter may help establish connections between genes and outcomes. Although a significant ExWAS finding does not necessarily imply causality, it provides an additional line of evidence that may corroborate MR and colocalization results. The quality control of whole-exome sequencing, creation of collapsing models, and methods used for ExWAS in the UK Biobank have been described previously[78]. Collapsing analyses were restricted to European ancestry individuals. For each gene, 11 collapsing models were constructed, each containing rare variants meeting specific inclusion criteria based on allele frequency as well as predicted effect[78]. Results were downloaded from the AstraZeneca PheWAS Portal (https://azphewas.com/)[78] for DBP (N = 395,230), SBP (N = 395,226), HbA1c (N = 400,339), LDL (N = 399,509) and Tg (N = 399,959), respectively. For each gene and each trait, we retained the collapsing model that showed the most significant association. Associations with a nominal p-value < 1.0x10^-5^ in collapsing analyses were considered suggestive.

### 5. Identification of known drug targets

We retrieved all drugs documented in the DrugBank database[79] that had reached Phase 4 of drug development, were marketed, and had not been cancelled by December 1, 2023. For each protein-trait association that had an FDR < 0.05 in primary MR analyses, we investigated whether the protein is a target, enzyme, carrier, or transporter of known therapies with indications relevant for the five cardiometabolic traits. These proteins are referred to as drug targets in this work. Specifically, for drugs that were indicated to treat hypertension, we examined whether colocalization evidence supported drug target-DBP or drug target-SBP associations. For drugs indicated to treat diabetes, we examined whether colocalization evidence supported drug target-HbA1c associations. For drugs indicated to treat hypercholesterolemia, we examined whether colocalization evidence supported drug target-LDL or drug target-Tg associations. Additionally, for drugs whose indications included myocardial infarction, heart failure, atrial fibrillation, or stroke, we examined whether colocalization evidence supported any associations between the drug targets and the five traits.

### 6. Colocalization based on meta-analyses of GWAS

For all protein-trait associations involving DBP, SBP, LDL, or Tg that were exclusively supported by one colocalization method, we reassessed colocalization evidence using large European ancestry-specific GWAS meta-analyses that included multiple cohorts and substantially larger sample sizes than the UK Biobank alone. Specifically, for DBP and SBP, we used summary statistics from Keaton et al., which included up to 1,028,980 individuals with measurements of blood pressure traits[80]. For LDL and Tg, we used summary statistics from Graham et al., which included up to 1,320,016 individuals with blood lipid measurements[63]. Details of these studies have been described previously[63,80]. We utilized the same LD reference panel based on 30,000 randomly selected, unrelated European ancestry individuals from the UK Biobank.

### 7. Additional evaluation of horizontal pleiotropy

For all protein-trait associations exclusively supported by one colocalization method, we conducted additional evaluation of horizontal pleiotropy, which may bias MR estimates. We obtained functional annotations for all genetic instruments used for these proteins from the Open Targets Genetics database[81,82], including the variant-to-gene (V2G) scores. The V2G scores were derived from an XGBoost model trained to quantify the functional connections between a variant and its neighboring genes[83]. A genetic instrument was deemed subject to a high risk of horizontal pleiotropy if the variant is located in the genic region of a gene other than the protein-coding gene, if it has been associated with the circulating level of another protein, or if it exhibits a stronger functional connection to a neighboring gene than the protein-coding gene based on V2G scores. If a genetic variant did not exhibit a high risk but has been associated with the expression level or alternative splicing of a neighboring gene other than the protein-coding gene, it would be considered to have a mild risk of horizontal pleiotropy. Additionally, for each instrument, we searched the NHGRI-EBI GWAS Catalog[84] to identify associations with any other human traits or diseases beyond circulating protein levels, retaining those reaching genome-wide significance (p-value < 5x10^-8^). However, a significant association in the GWAS Catalog does not necessarily indicate that the variant is subject to horizontal pleiotropy, as it may instead reflect pleiotropic effects of the protein.

### 8. Phenotypic impact in mouse mutants

We assessed the biological relevance of genes involved in protein-trait associations exclusively supported by one colocalization method. Phenotypic changes and abnormalities in mouse mutants were retrieved from the Mouse Genome Informatics database[85] for homologous genes in mice. Moreover, characterizations of single-gene knockout mice in comparison to wild types were retrieved from the International Mouse Phenotyping Consortium database[44], where phenotypic changes with a nominal p-value < 1.0x10^-4^ were considered suggestive.

## Code availability

coloc and coloc+SuSiE are available at https://github.com/chr1swallace/coloc.

PWCoCo is available at https://github.com/jwr-git/pwcoco.

SharePro is available at https://github.com/zhwm/SharePro_coloc.

All processed results are included in this manuscript. Raw colocalization results are available in the Figshare repository: https://figshare.com/s/f2ccd417c42a04a4f9be.

Computational scripts are available in the BioCode repository: https://ngdc.cncb.ac.cn/biocode/ (accession number BT007988).

## CRediT author statement

Wenmin Zhang: Investigation, Methodology, Resources, Formal analysis, Writing – original draft, Writing – review & editing

Satoshi Yoshiji: Resources, Writing – review & editing

Robert Sladek: Investigation, Writing – review & editing

Josée Dupuis: Investigation, Resources, Writing – review & editing

Tianyuan Lu: Conceptualization, Investigation, Methodology, Resources, Formal analysis, Writing – original draft, Writing – review & editing

## Competing interest

W.Z. was the developer of SharePro, one of the colocalization methods under investigation. R.S., J.D. and T.L. co-authored the publication describing SharePro. T.L. and W.Z. have been consulting for Five Prime Sciences Inc. The research presented in this paper was conducted independently, and 5 Prime Sciences Inc. was not involved in the design, execution, analysis, or interpretation of the study. The other authors declare no conflicts of interest.

## Acknowledgements

This study was enabled, in part, by support from Calcul Québec and Digital Research Alliance of Canada. W.Z. has been supported by a Postdoctoral Fellowship from the Institut de valorisation des données (IVADO), funded by the Canada First Research Excellence Fund. T.L. has been supported by start-up funding from the Office of the Vice Chancellor for Research and Graduate Education, School of Medicine and Public Health, and Department of Population Health Sciences at the University of Wisconsin-Madison. The funders have no role in study design; collection, management, analysis and interpretation of data; or the decision to submit for publication.

## Supplementary material

**Figure S1. Manhattan plot showing genome-wide distribution of associations between circulating protein levels and diastolic blood pressure.**

Dots present the coding-genes of circulating proteins under investigation. The dashed line indicates the Bonferroni-corrected significance threshold. The dash-dotted line indicates the false discovery rate threshold of 0.05. Full test statistics of Mendelian randomization are available in Supplementary Table S2.

**Figure S2. Manhattan plot showing genome-wide distribution of associations between circulating protein levels and systolic blood pressure.**

Dots present the coding-genes of circulating proteins under investigation. The dashed line indicates the Bonferroni-corrected significance threshold. The dash-dotted line indicates the false discovery rate threshold of 0.05. Full test statistics of Mendelian randomization are available in Supplementary Table S2.

**Figure S3. Manhattan plot showing genome-wide distribution of associations between circulating protein levels and hemoglobin A1c.**

Dots present the coding-genes of circulating proteins under investigation. The dashed line indicates the Bonferroni-corrected significance threshold. The dash-dotted line indicates the false discovery rate threshold of 0.05. Full test statistics of Mendelian randomization are available in Supplementary Table S2.

**Figure S4. Manhattan plot showing genome-wide distribution of associations between circulating protein levels and low-density lipoprotein cholesterol.**

Dots present the coding-genes of circulating proteins under investigation. The dashed line indicates the Bonferroni-corrected significance threshold. The dash-dotted line indicates the false discovery rate threshold of 0.05. Full test statistics of Mendelian randomization are available in Supplementary Table S2.

**Figure S5. Manhattan plot showing genome-wide distribution of associations between circulating protein levels and triglycerides.**

Dots present the coding-genes of circulating proteins under investigation. The dashed line indicates the Bonferroni-corrected significance threshold. The dash-dotted line indicates the false discovery rate threshold of 0.05. Full test statistics of Mendelian randomization are available in Supplementary Table S2.

**Figure S6. Illustration of genetic associations with circulating UBASH3B level and with HbA1c.** The lead variants of cis-pQTLs used as genetic instruments are indicated. Genetic variants located in a ±500kb window centered around each genetic instrument are plotted with their significance in respective studies, and colored by the magnitude of correlation (linkage disequilibrium r^2^) with the corresponding instrument. Two horizontal dashed lines and two vertical dashed lines represent p-value thresholds of 1.0x10^-5^ and 5.0x10^-8^, respectively. Protein names include SOMAmer identifiers. Cis-pQTL: cis-protein quantitative trait locus.

**Figure S7. Comparison of colocalization probabilities inferred using four Bayesian colocalization methods with the default prior.**

Mendelian randomization-identified associations with a false discovery rate < 0.5 are included.

**Figure S8. Comparison of colocalization evidence and significance of associations in Mendelian randomization.**

Dots represent protein-trait associations with a false discovery rate < 0.5, colored with respect to colocalization evidence generated by coloc. The vertical dashed line represents the Bonferroni-corrected significance threshold. Two horizontal dashed lines represent colocalization probability thresholds of 0.5 and 0.8, respectively.

**Figure S9. Comparison of colocalization evidence and significance of associations in Mendelian randomization.**

Dots represent protein-trait associations with a false discovery rate < 0.5, colored with respect to colocalization evidence generated by coloc+SuSiE. The vertical dashed line represents the Bonferroni-corrected significance threshold. Two horizontal dashed lines represent colocalization probability thresholds of 0.5 and 0.8, respectively.

**Figure S10. Comparison of colocalization evidence and significance of associations in Mendelian randomization.**

Dots represent protein-trait associations with a false discovery rate < 0.5, colored with respect to colocalization evidence generated by PWCoCo. The vertical dashed line represents the Bonferroni-corrected significance threshold. Two horizontal dashed lines represent colocalization probability thresholds of 0.5 and 0.8, respectively.

**Figure S11. Comparison of colocalization evidence and significance of associations in Mendelian randomization.**

Dots represent protein-trait associations with a false discovery rate < 0.5, colored with respect to colocalization evidence generated by SharePro. The vertical dashed line represents the Bonferroni-corrected significance threshold. Two horizontal dashed lines represent colocalization probability thresholds of 0.5 and 0.8, respectively.

**Figure S12. Prior sensitivity analyses for diastolic blood pressure.**

For protein-trait associations supported by strong colocalization evidence using coloc, coloc+SuSiE, PWCoCo, and SharePro with the default prior of 1.0x10^-5^, respectively, the proportions of these associations supported by each method using different prior settings are summarized.

**Figure S13. Prior sensitivity analyses for systolic blood pressure.**

For protein-trait associations supported by strong colocalization evidence using coloc, coloc+SuSiE, PWCoCo, and SharePro with the default prior of 1.0x10^-5^, respectively, the proportions of these associations supported by each method using different prior settings are summarized.

**Figure S14. Prior sensitivity analyses for hemoglobin A1c.**

For protein-trait associations supported by strong colocalization evidence using coloc, coloc+SuSiE, PWCoCo, and SharePro with the default prior of 1.0x10^-5^, respectively, the proportions of these associations supported by each method using different prior settings are summarized.

**Figure S15. Prior sensitivity analyses for low-density lipoprotein cholesterol.**

For protein-trait associations supported by strong colocalization evidence using coloc, coloc+SuSiE, PWCoCo, and SharePro with the default prior of 1.0x10^-5^, respectively, the proportions of these associations supported by each method using different prior settings are summarized.

**Figure S16. Prior sensitivity analyses for triglycerides.**

For protein-trait associations supported by strong colocalization evidence using coloc, coloc+SuSiE, PWCoCo, and SharePro with the default prior of 1.0x10^-5^, respectively, the proportions of these associations supported by each method using different prior settings are summarized.

**Figure S17. Prior sensitivity analyses for associations involving significant findings in exome-wide association studies.**

For each protein-trait association, colocalization probabilities inferred with different prior settings are indicated for each method. The grey dashed lines indicate colocalization probabilities of 0.8 and 0.5, respectively.

**Figure S18. Prior sensitivity analyses for associations involving successful drug targets.**

For each protein-trait association, colocalization probabilities inferred with different prior settings are indicated for each method. The grey dashed lines indicate colocalization probabilities of 0.8 and 0.5, respectively.

**Figure S19. Prior sensitivity analyses for associations exclusively supported by SharePro.**

For each protein-trait association, colocalization probabilities inferred with different prior settings are indicated for each method. The grey dashed lines indicate colocalization probabilities of 0.8 and 0.5, respectively.

**Table S1. Association results for all instruments.**

**Table S2. Mendelian randomization results.**

Beta represents the change of standardized outcome per 1 SD increase in genetically predicted circulating protein abundance.

**Table S3. Assessment of colocalization for protein-trait associations with a p-value > 0.9 in Mendelian randomization.**

**Table S4. Assessment of colocalization for significant protein-trait associations with a varying prior colocalization probability.**

**Table S5. Comparison of runtime and memory usage across all analyzed protein-trait associations.**

All analyses were performed using a single 2 x Intel E5-2683 v4 Broadwell @ 2.1GHz CPU core on Compute Canada infrastructure.

**Table S6. Findings from exome-wide association studies.**

Summary statistics for the most significant collapsing model of each gene are shown. Genes that have no collapsing model with a linear regression p-value < 0.1 are not included.

**Table S7. Known drug targets in DrugBank amongst protein-trait associations with a false discovery rate < 0.5.**

The drugs that have been approved and marketed, and have not been cancelled are included.

**Table S8. Assessment of colocalization for associations exclusively supported by SharePro based on additional GWAS.**

Colocalization probability was obtained using SharePro with the default prior of 1x10^-5^. Table S9. Open Targets Genetics annotation of genetic instruments.

**Table S10. Evaluation of risk of horizontal pleiotropy.**

**Table S11. Known associations between instruments and human traits and diseases. Table S12. Mouse mutants and suggestive phenotypic changes.**

## Notes

### Summary of Updates

This version has expanded discussion of the findings. The results remain unchanged.

