## Supplementary Figures for "Benchmarking Bayesian colocalization methods in validating Mendelian randomization-identified targets"

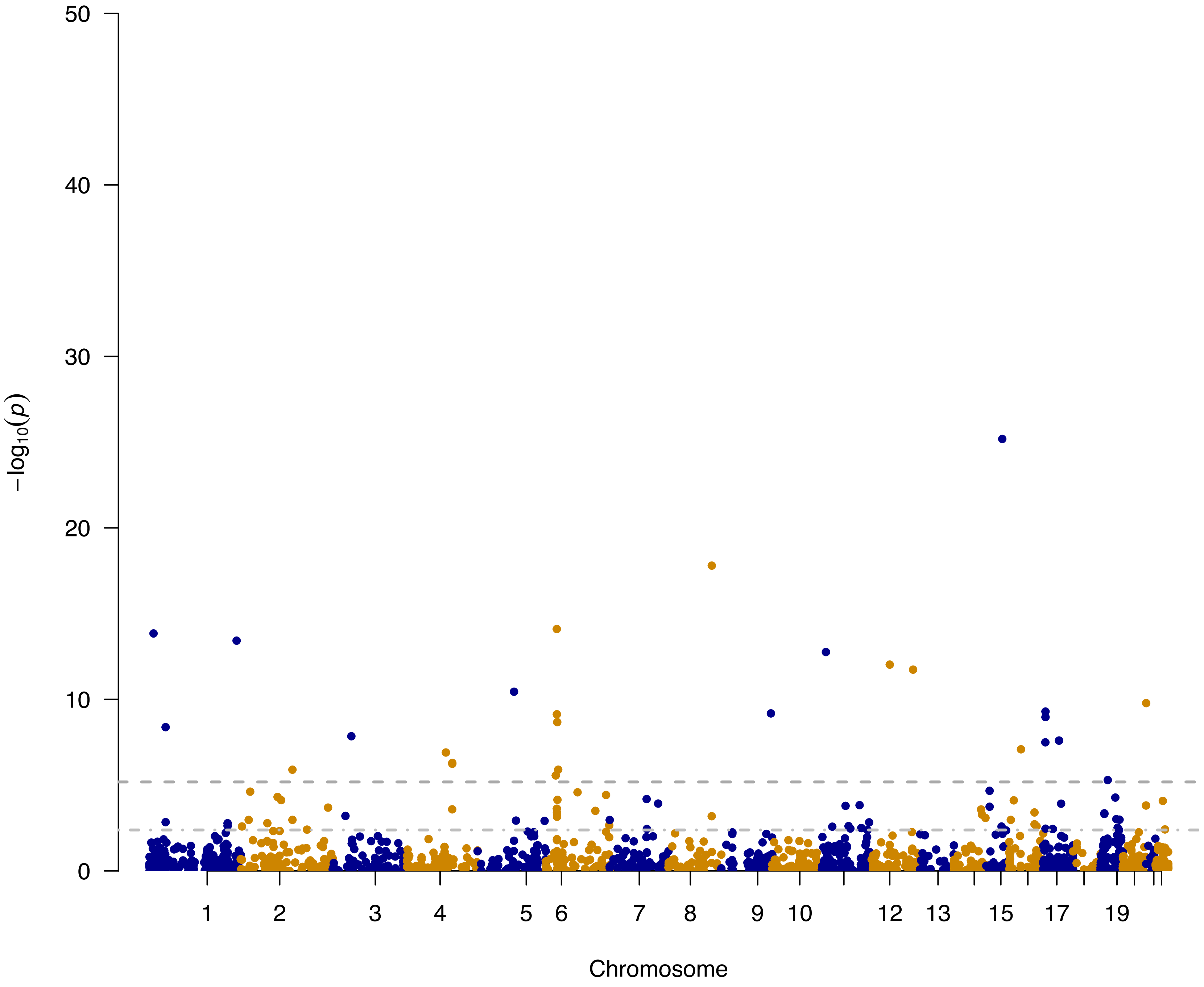


**Figure S1.** Manhattan plot showing genome-wide distribution of associations between circulating protein levels and diastolic blood pressure. Dots present the coding-genes of circulating proteins under investigation. The dashed line indicates the Bonferroni-corrected significance threshold. The dash-dotted line indicates the false discovery rate threshold of 0.05. Full test statistics of Mendelian randomization are available in Supplementary Table S2.

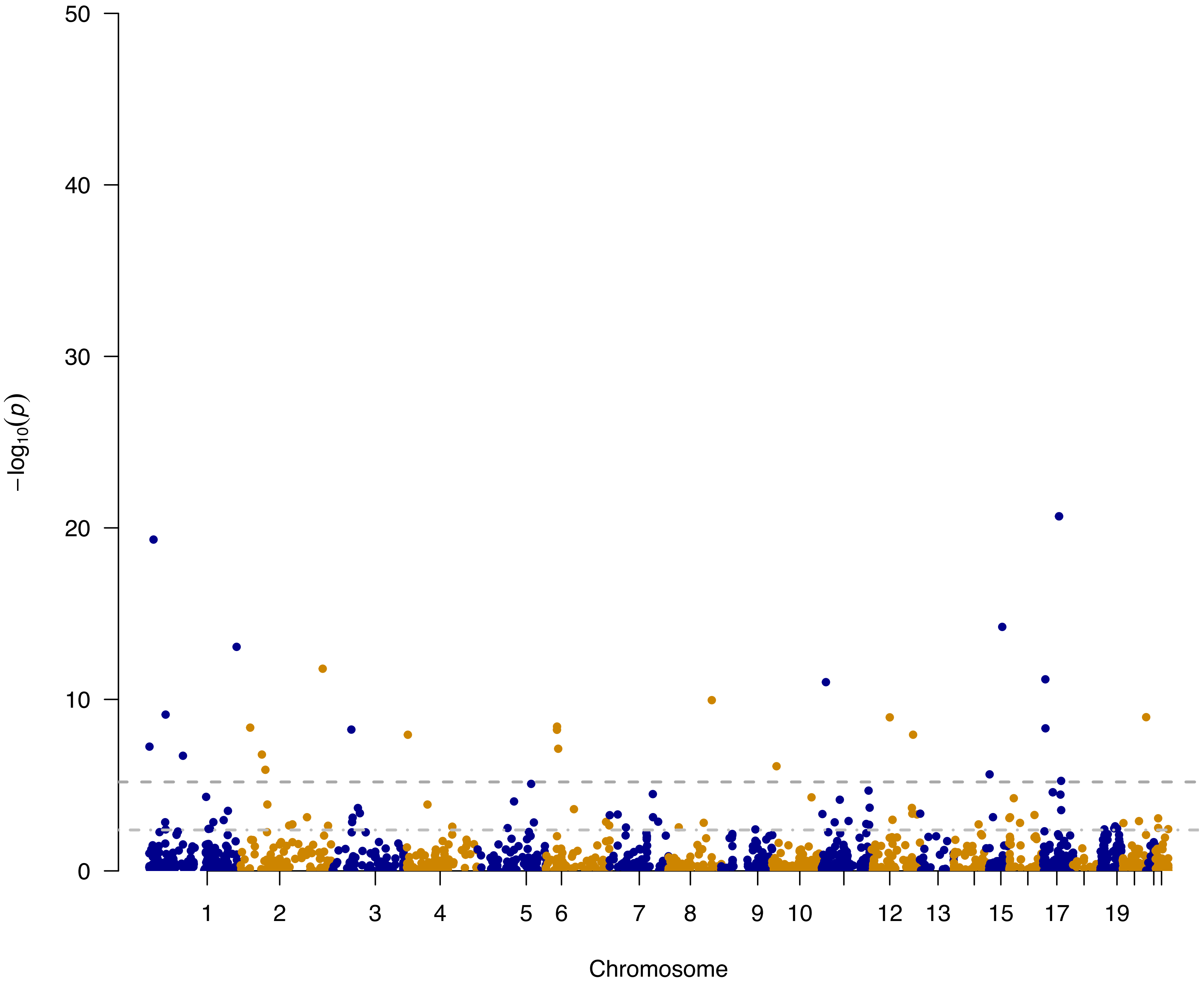


**Figure S2.** Manhattan plot showing genome-wide distribution of associations between circulating protein levels and systolic blood pressure. Dots present the coding-genes of circulating proteins under investigation. The dashed line indicates the Bonferroni-corrected significance threshold. The dash-dotted line indicates the false discovery rate threshold of 0.05. Full test statistics of Mendelian randomization are available in Supplementary Table S2.

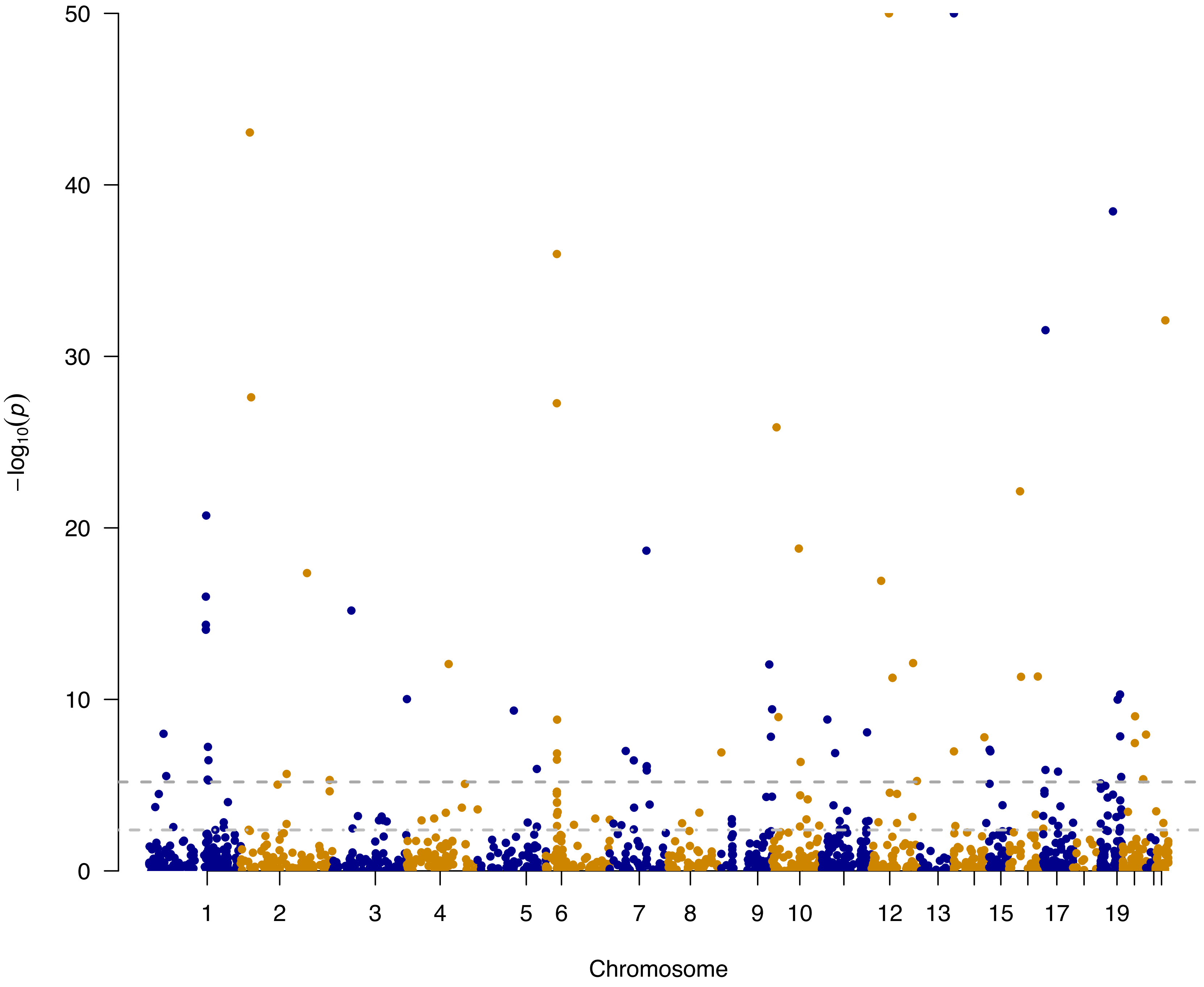


**Figure S3.** Manhattan plot showing genome-wide distribution of associations between circulating protein levels and hemoglobin A1c. Dots present the coding-genes of circulating proteins under investigation. The dashed line indicates the Bonferroni-corrected significance threshold. The dash-dotted line indicates the false discovery rate threshold of 0.05. Full test statistics of Mendelian randomization are available in Supplementary Table S2.

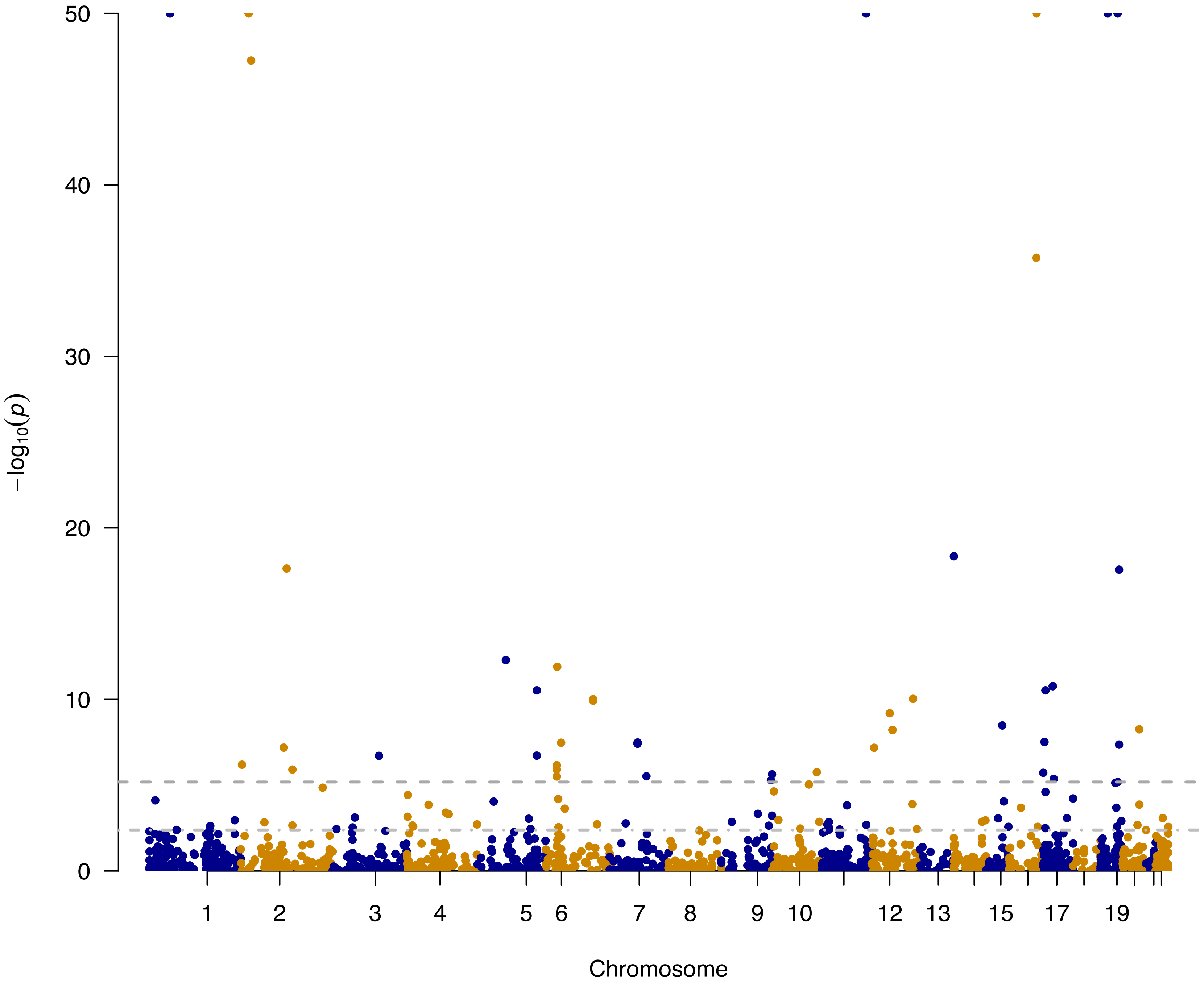


**Figure S4.** Manhattan plot showing genome-wide distribution of associations between circulating protein levels and low-density lipoprotein cholesterol. Dots present the coding-genes of circulating proteins under investigation. The dashed line indicates the Bonferroni-corrected significance threshold. The dash-dotted line indicates the false discovery rate threshold of 0.05. Full test statistics of Mendelian randomization are available in Supplementary Table S2.

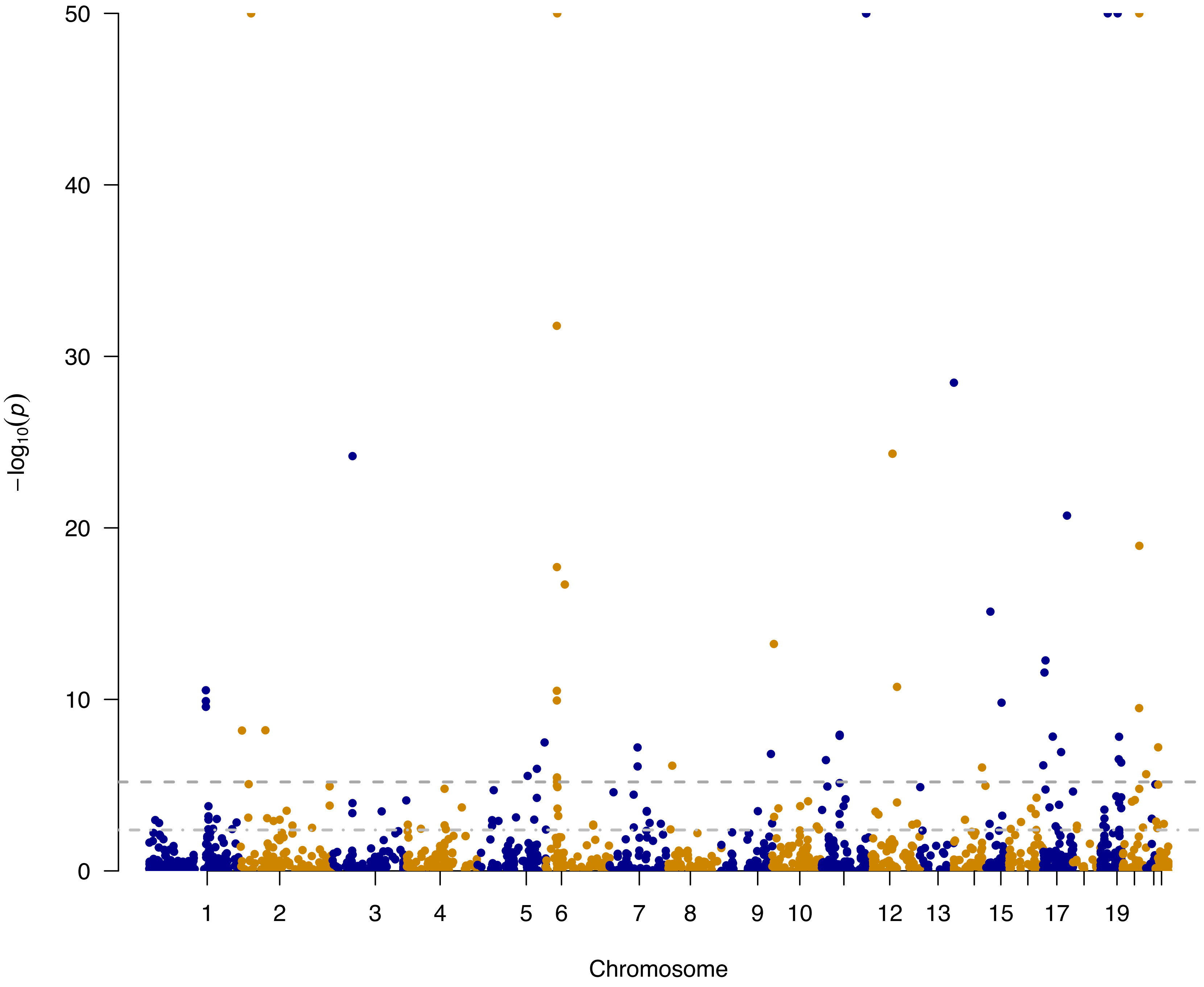


**Figure S5.** Manhattan plot showing genome-wide distribution of associations between circulating protein levels and triglycerides. Dots present the coding-genes of circulating proteins under investigation. The dashed line indicates the Bonferroni-corrected significance threshold. The dash-dotted line indicates the false discovery rate threshold of 0.05. Full test statistics of Mendelian randomization are available in Supplementary Table S2.

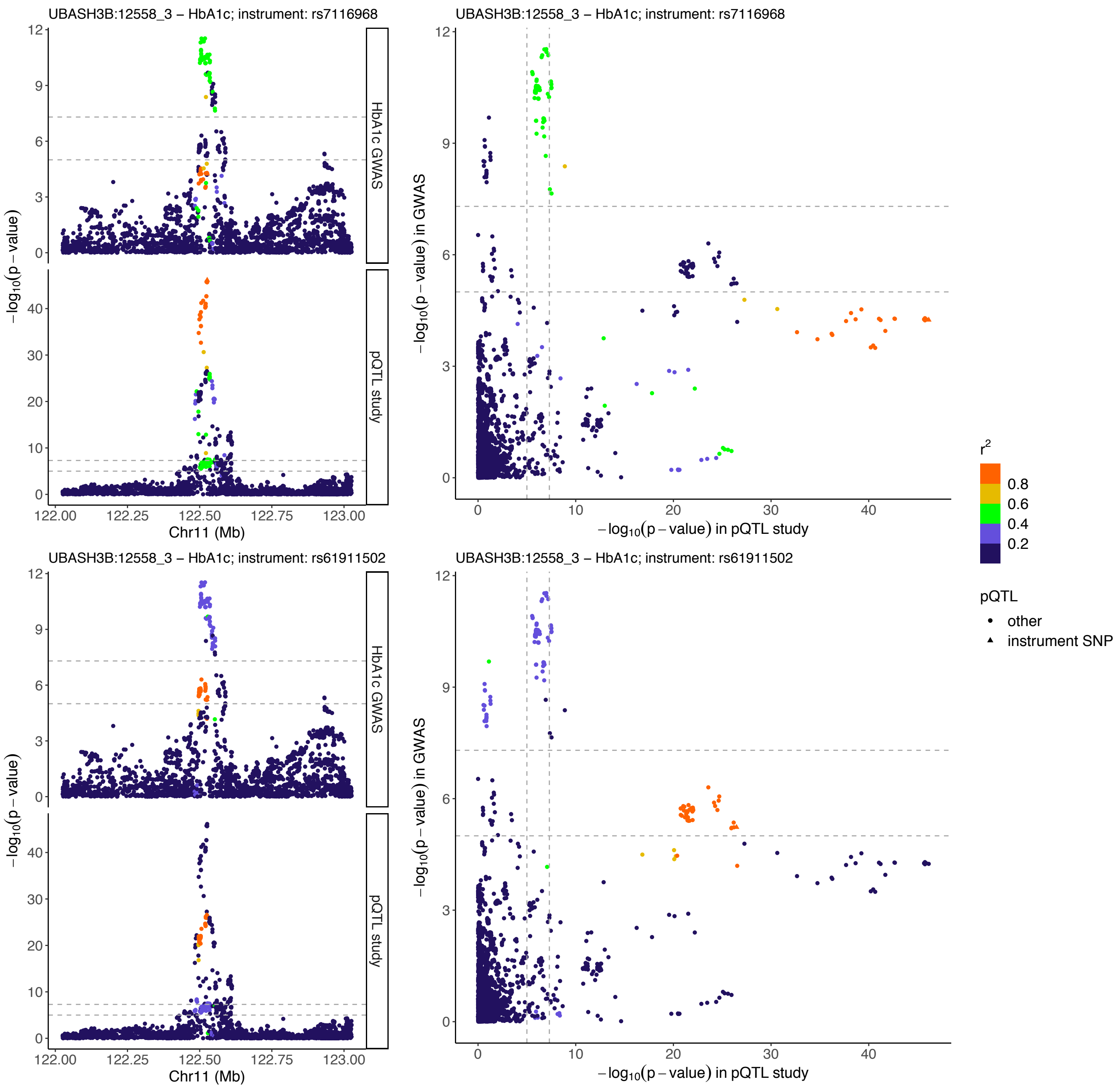


**Figure S6.** Illustration of genetic associations with circulating UBASH3B level and with HbA1c. The lead variants of cis-protein quantitative trait loci (cis-pQTLs) used as genetic instruments are indicated. Genetic variants located in a ±500kb window centered around each genetic instrument are plotted with their significance in respective studies, and colored by the magnitude of correlation (linkage disequilibrium $r^2$) with the corresponding instrument. Two horizontal dashed lines and two vertical dashed lines represent p-value thresholds of $1.0 \times 10^{-5}$ and $5.0 \times 10^{-8}$, respectively. Protein names include SOMAmer identifiers.

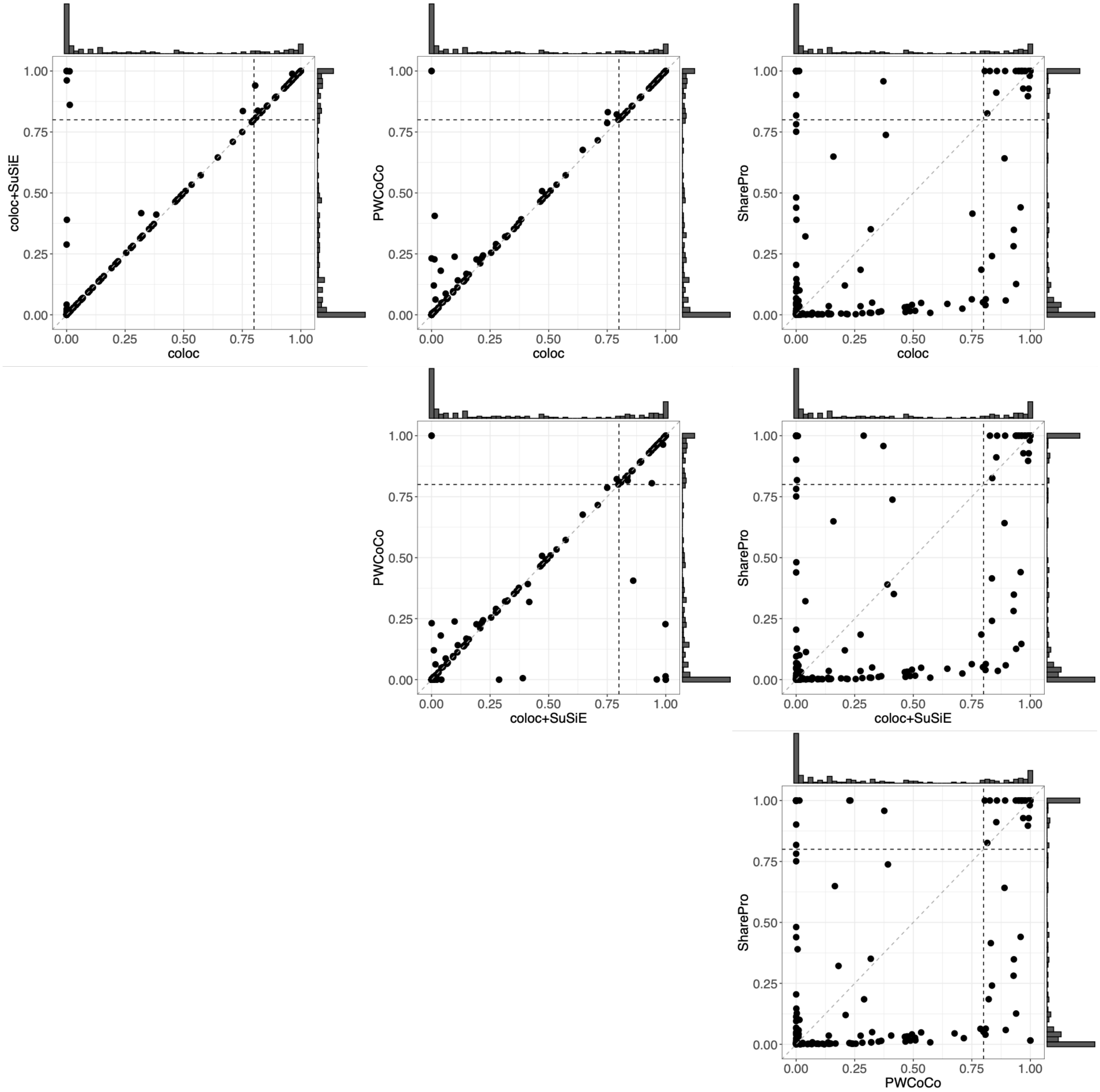


**Figure S7.** Comparison of colocalization probabilities inferred using four Bayesian colocalization methods with the default prior. Mendelian randomization-identified associations with a false discovery rate < 0.5 are included.

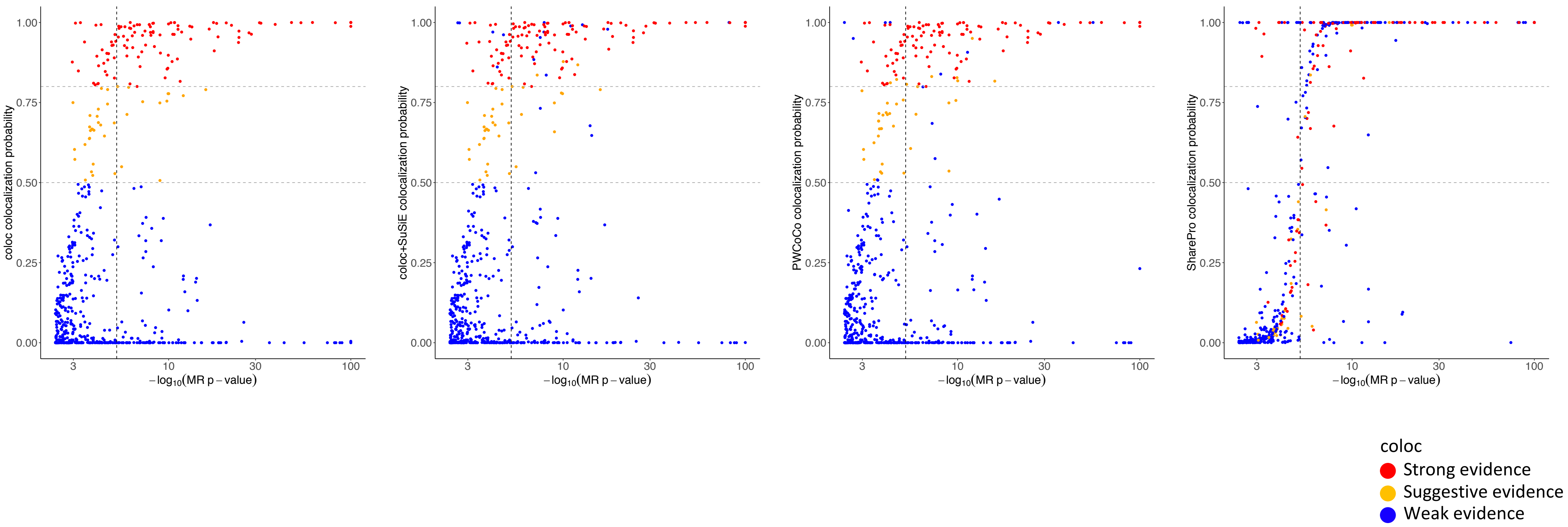


**Figure S8.** Comparison of colocalization evidence and significance of associations in Mendelian randomization. Dots represent protein-trait associations with a false discovery rate < 0.5, colored with respect to colocalization evidence generated by coloc. The vertical dashed line represents the Bonferroni-corrected significance threshold. Two horizontal dashed lines represent colocalization probability thresholds of 0.5 and 0.8, respectively.

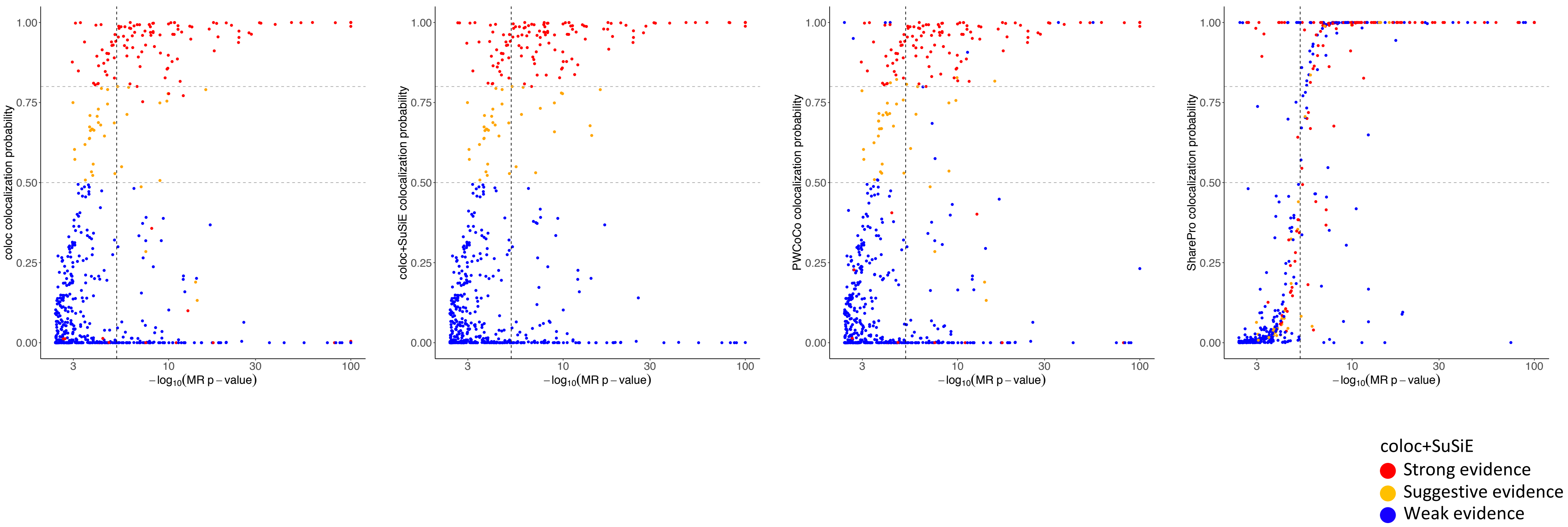


**Figure S9.** Comparison of colocalization evidence and significance of associations in Mendelian randomization. Dots represent protein-trait associations with a false discovery rate < 0.5, colored with respect to colocalization evidence generated by coloc+SuSiE. The vertical dashed line represents the Bonferroni-corrected significance threshold. Two horizontal dashed lines represent colocalization probability thresholds of 0.5 and 0.8, respectively.

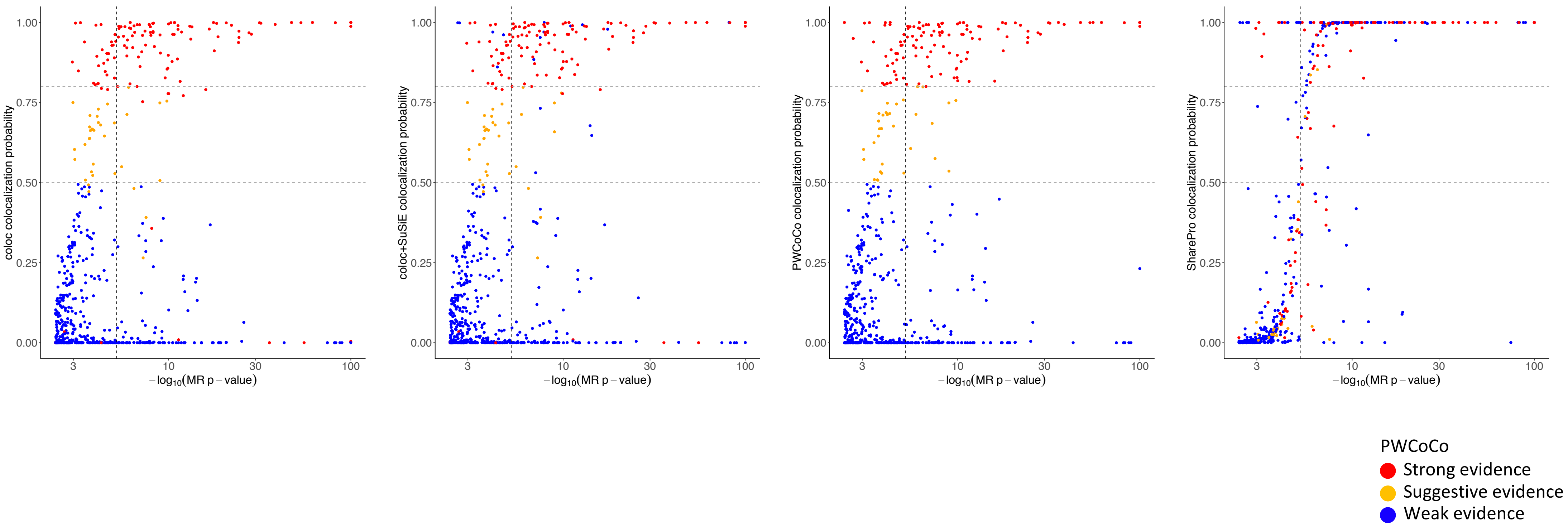


**Figure S10.** Comparison of colocalization evidence and significance of associations in Mendelian randomization. Dots represent protein-trait associations with a false discovery rate < 0.5, colored with respect to colocalization evidence generated by PWCoCo. The vertical dashed line represents the Bonferroni-corrected significance threshold. Two horizontal dashed lines represent colocalization probability thresholds of 0.5 and 0.8, respectively.

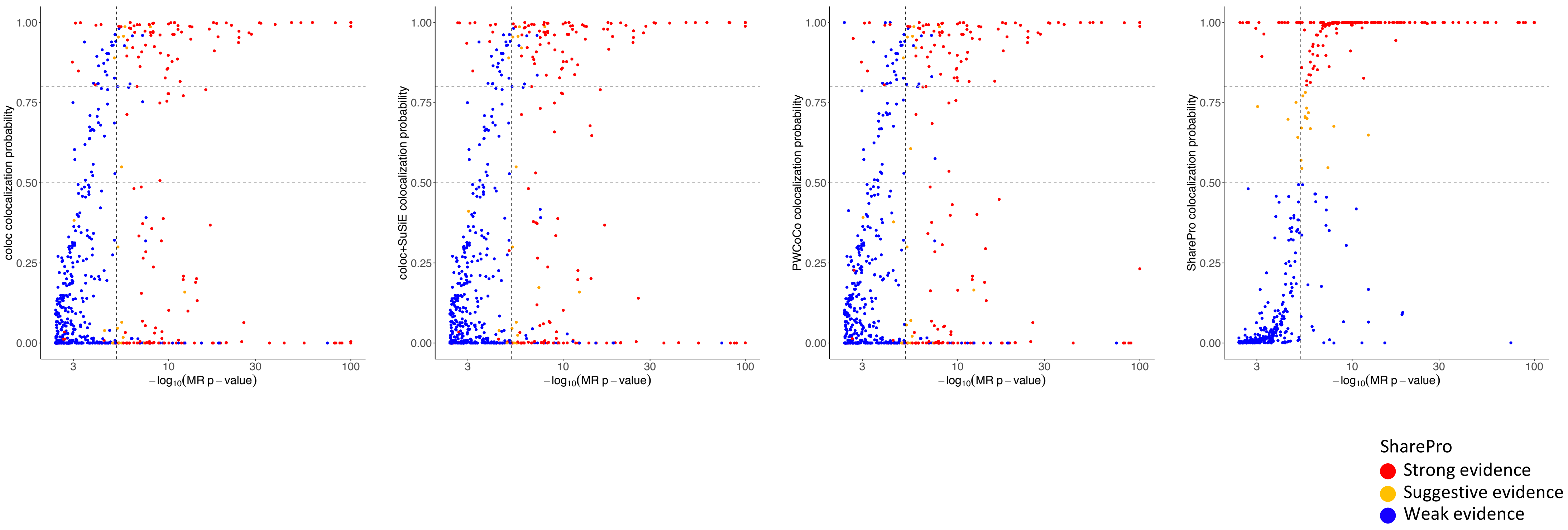


**Figure S11.** Comparison of colocalization evidence and significance of associations in Mendelian randomization. Dots represent protein-trait associations with a false discovery rate < 0.5, colored with respect to colocalization evidence generated by SharePro. The vertical dashed line represents the Bonferroni-corrected significance threshold. Two horizontal dashed lines represent colocalization probability thresholds of 0.5 and 0.8, respectively.

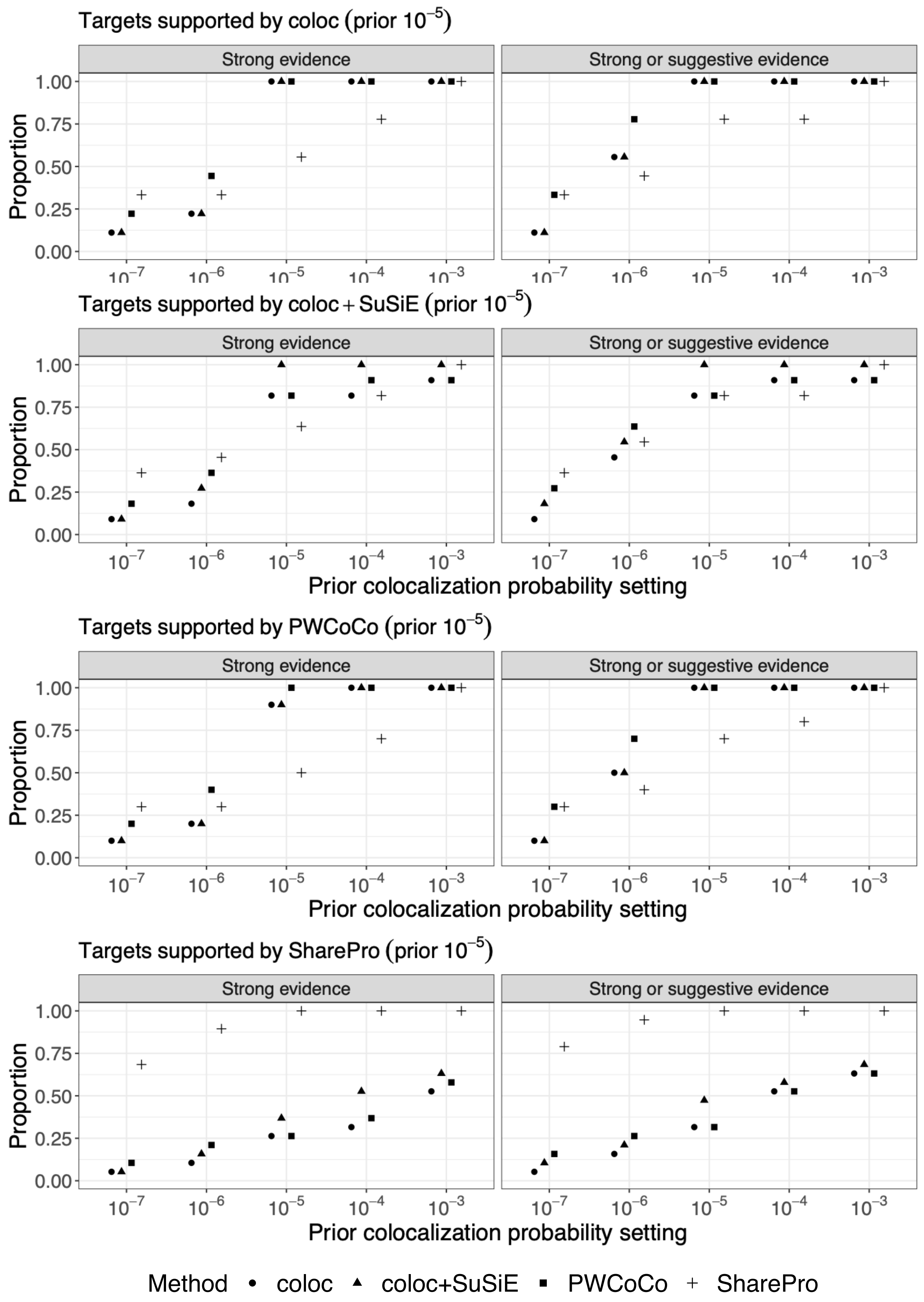


**Figure S12.** Prior sensitivity analyses for diastolic blood pressure. For protein-trait associations supported by strong colocalization evidence using coloc, coloc+SuSiE, PWCoCo, and SharePro with the default prior of $1.0 \times 10^{-5}$, respectively, the proportions of these associations supported by each method using different prior settings are summarized.

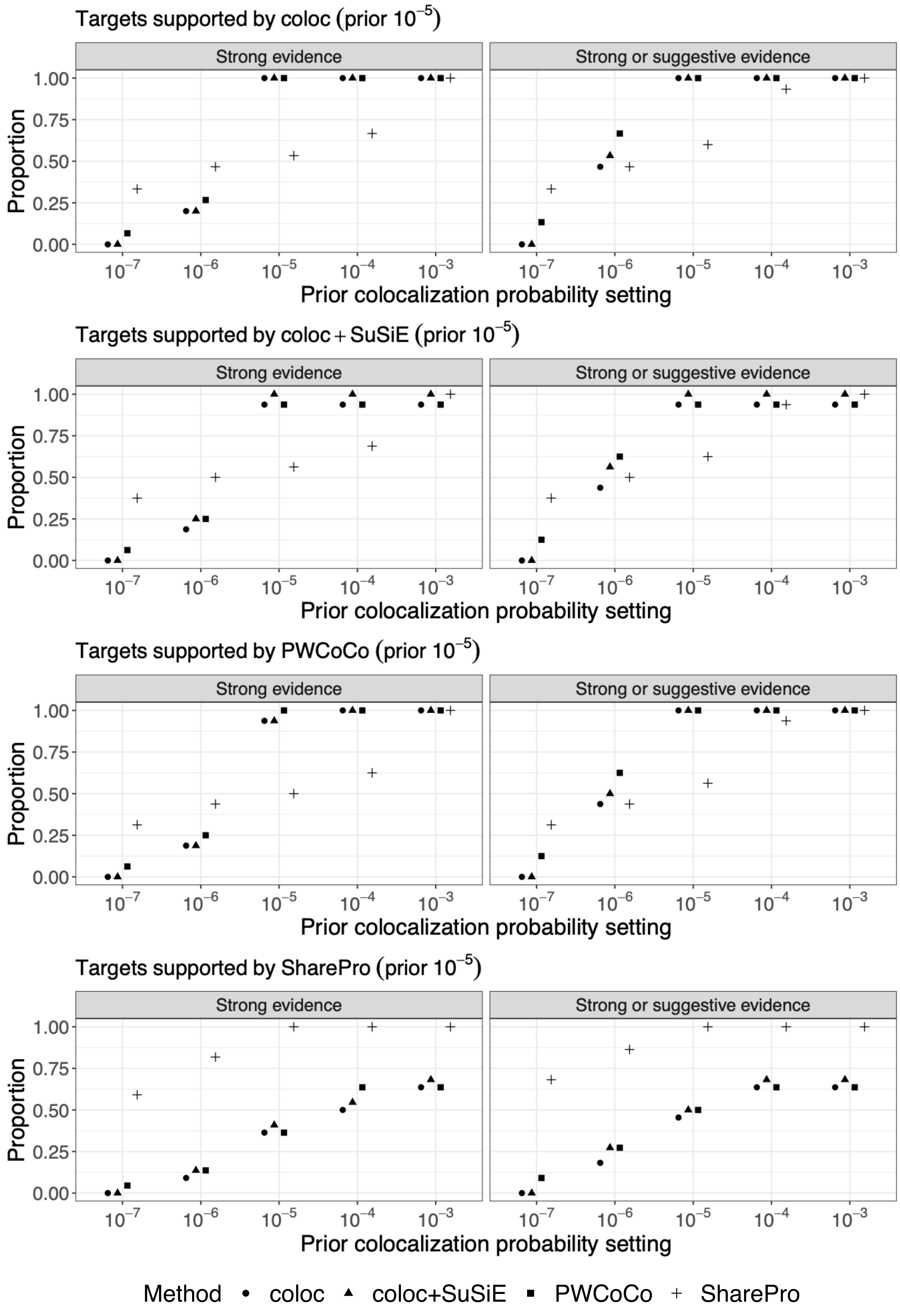


**Figure S13.** Prior sensitivity analyses for systolic blood pressure. For protein-trait associations supported by strong colocalization evidence using coloc, coloc+SuSiE, PWCoCo, and SharePro with the default prior of 1.0x$10^{-5}$, respectively, the proportions of these associations supported by each method using different prior settings are summarized.

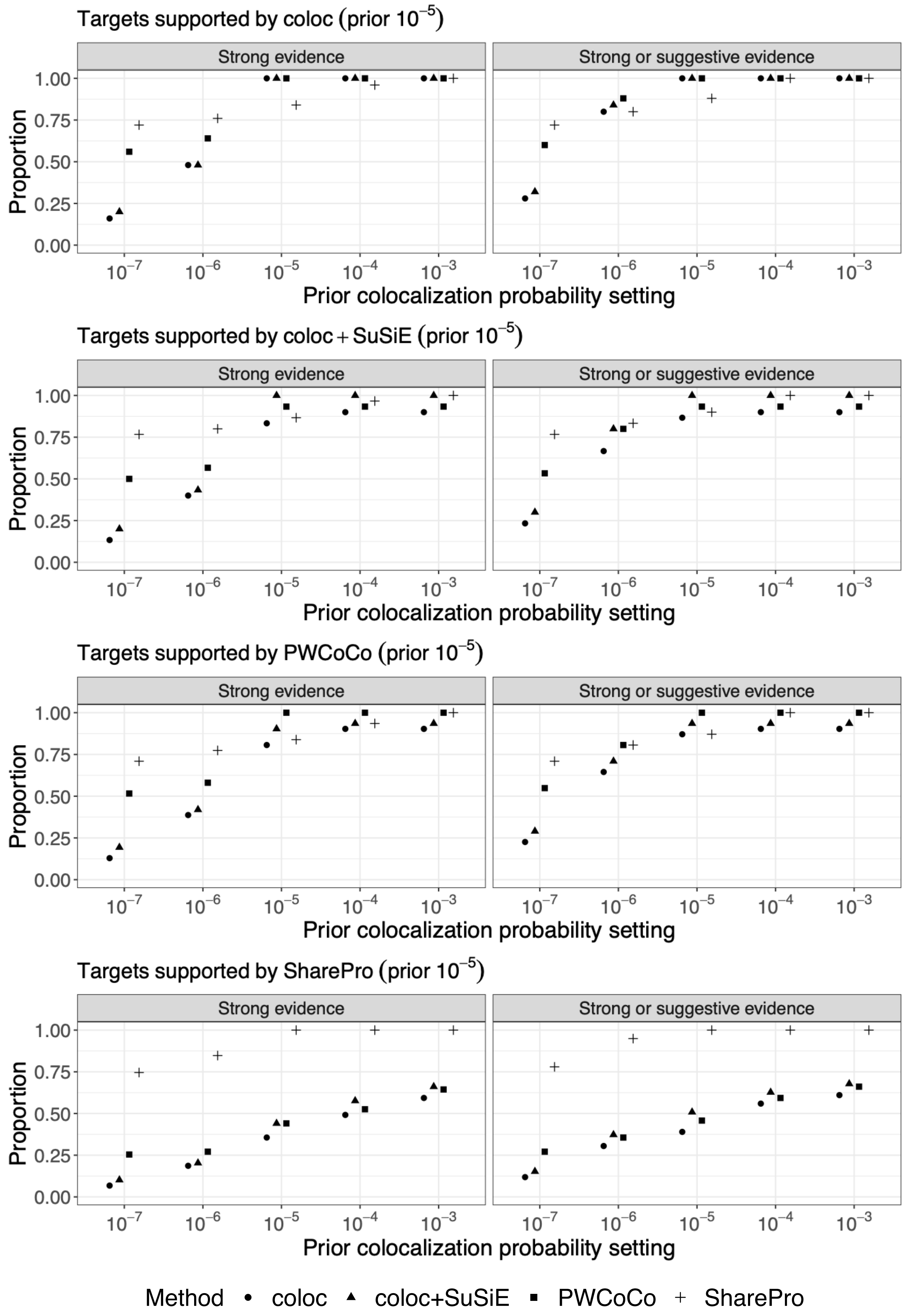


**Figure S14.** Prior sensitivity analyses for hemoglobin A1c. For protein-trait associations supported by strong colocalization evidence using coloc, coloc+SuSiE, PWCoCo, and SharePro with the default prior of $1.0x10^{-5}$, respectively, the proportions of these associations supported by each method using different prior settings are summarized.

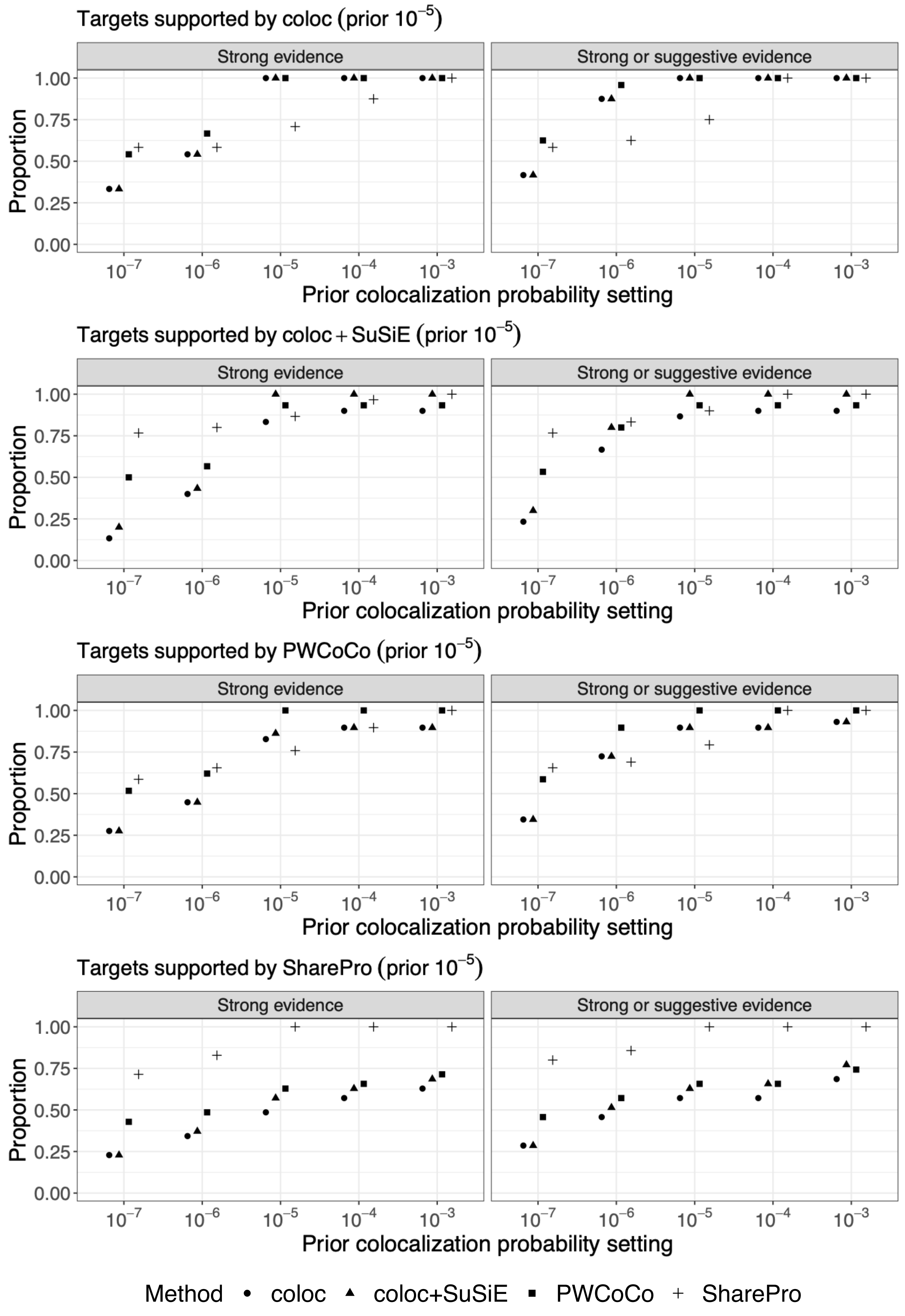


**Figure S15.** Prior sensitivity analyses for low-density lipoprotein cholesterol. For protein-trait associations supported by strong colocalization evidence using coloc, coloc+SuSiE, PWCoCo, and SharePro with the default prior of 1.0x10$^{-5}$, respectively, the proportions of these associations supported by each method using different prior settings are summarized.

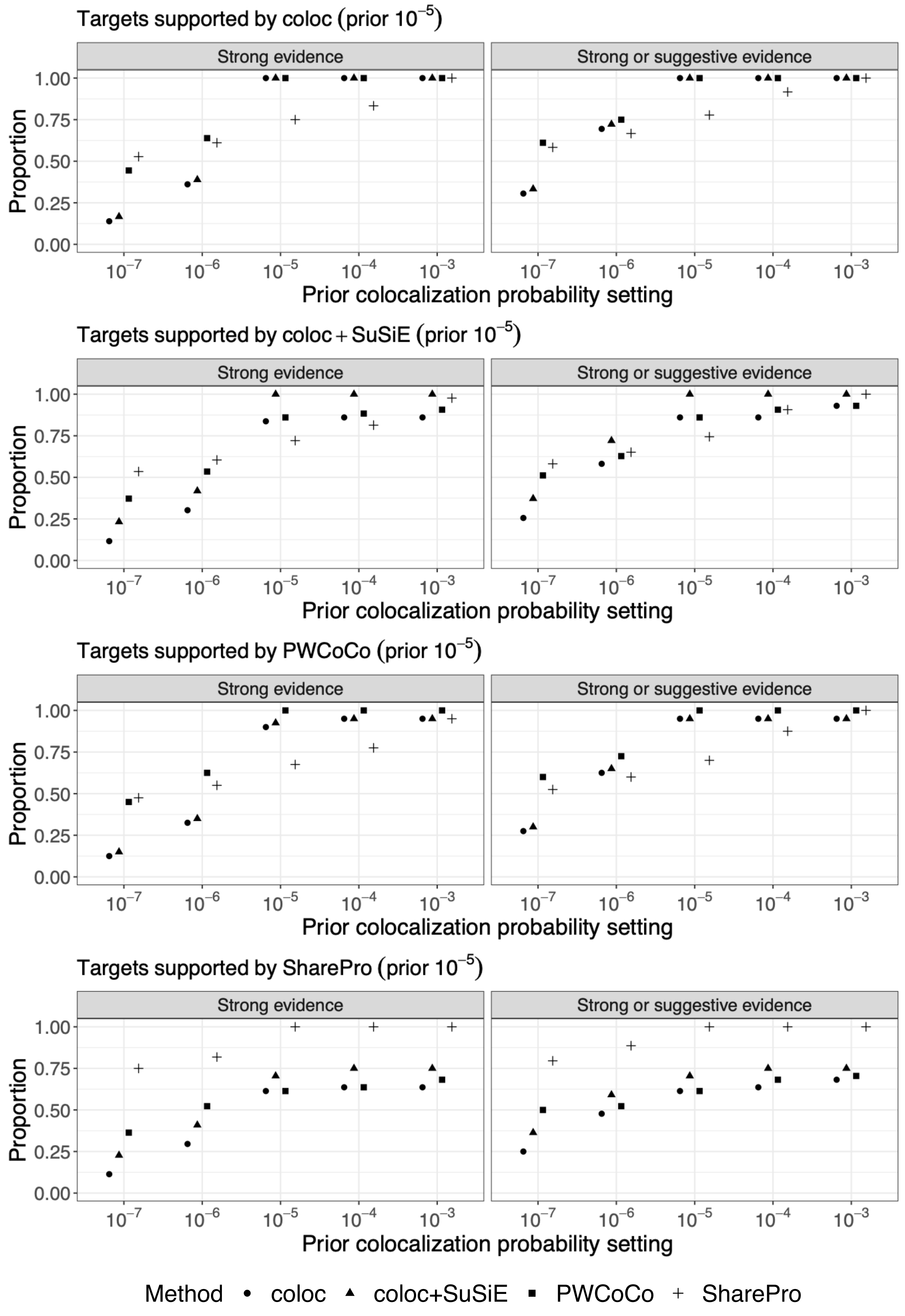


**Figure S16.** Prior sensitivity analyses for triglycerides. For protein-trait associations supported by strong colocalization evidence using coloc, coloc+SuSiE, PWCoCo, and SharePro with the default prior of $1.0 \times 10^{-5}$, respectively, the proportions of these associations supported by each method using different prior settings are summarized.

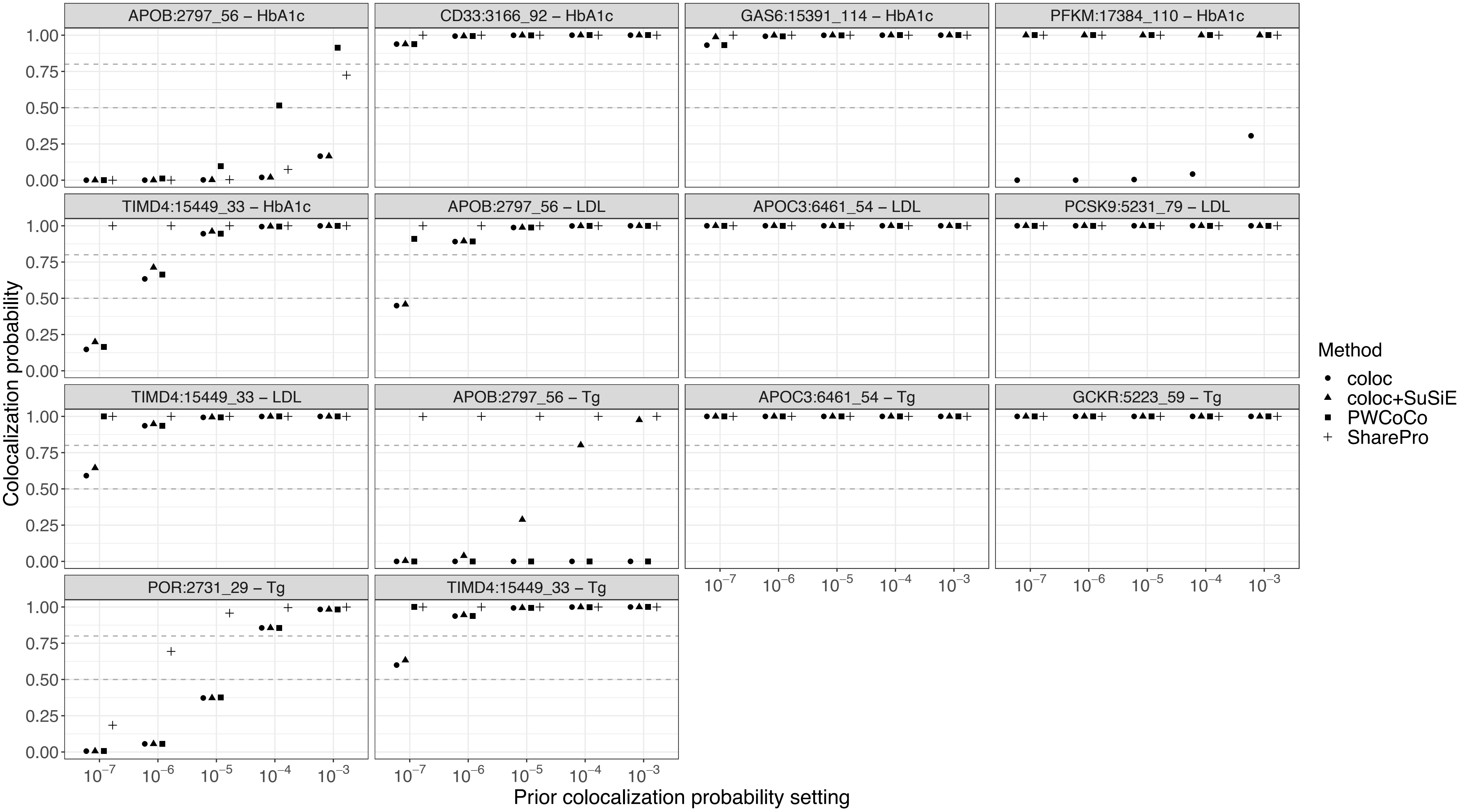


**Figure S17.** Prior sensitivity analyses for associations involving significant findings in exome-wide association studies. For each protein-trait association, colocalization probabilities inferred with different prior settings are indicated for each method. The grey dashed lines indicate colocalization probabilities of 0.8 and 0.5, respectively.

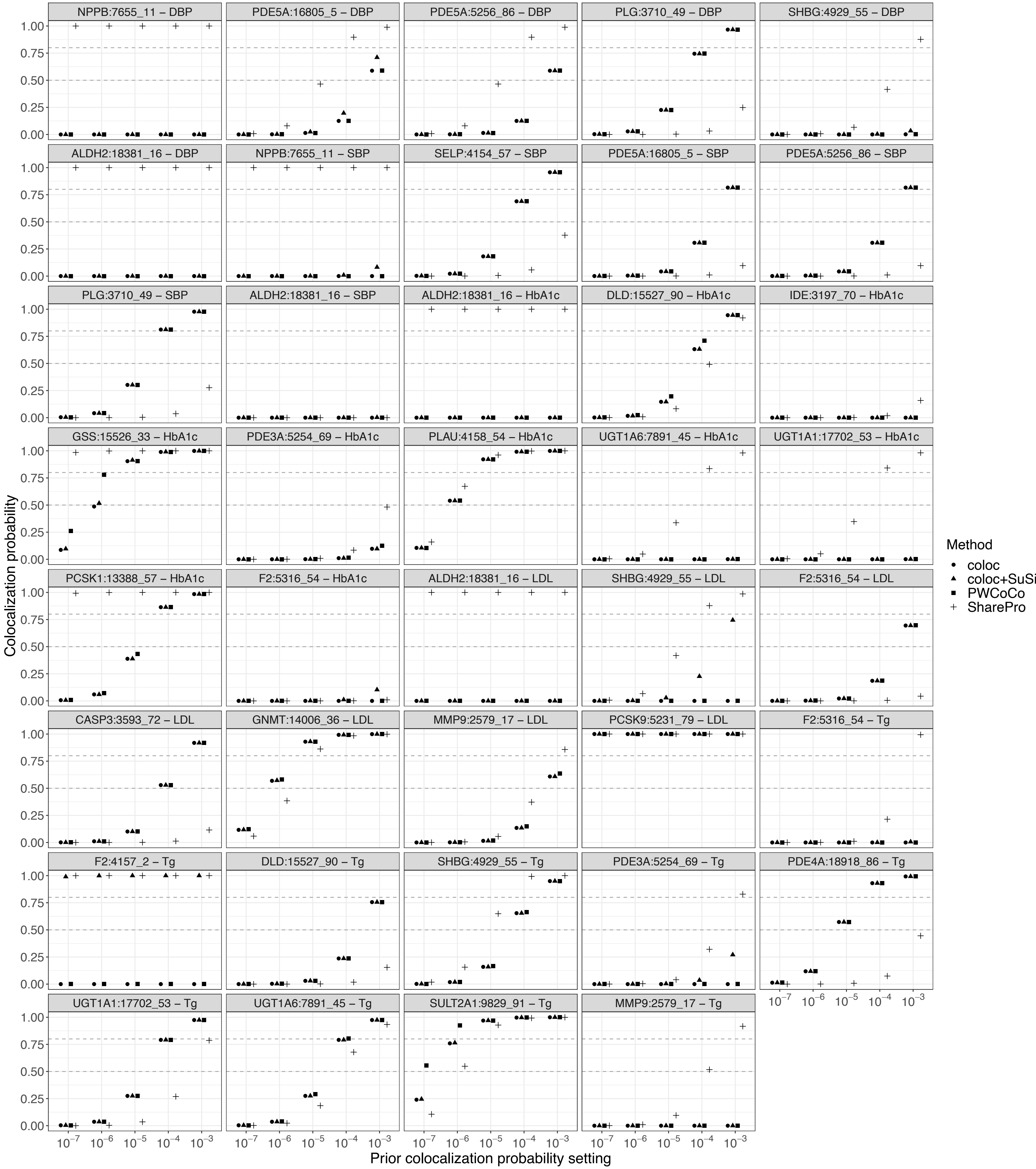


**Figure S18.** Prior sensitivity analyses for associations involving successful drug targets. For each protein-trait association, colocalization probabilities inferred with different prior settings are indicated for each method. The grey dashed lines indicate colocalization probabilities of 0.8 and 0.5, respectively.

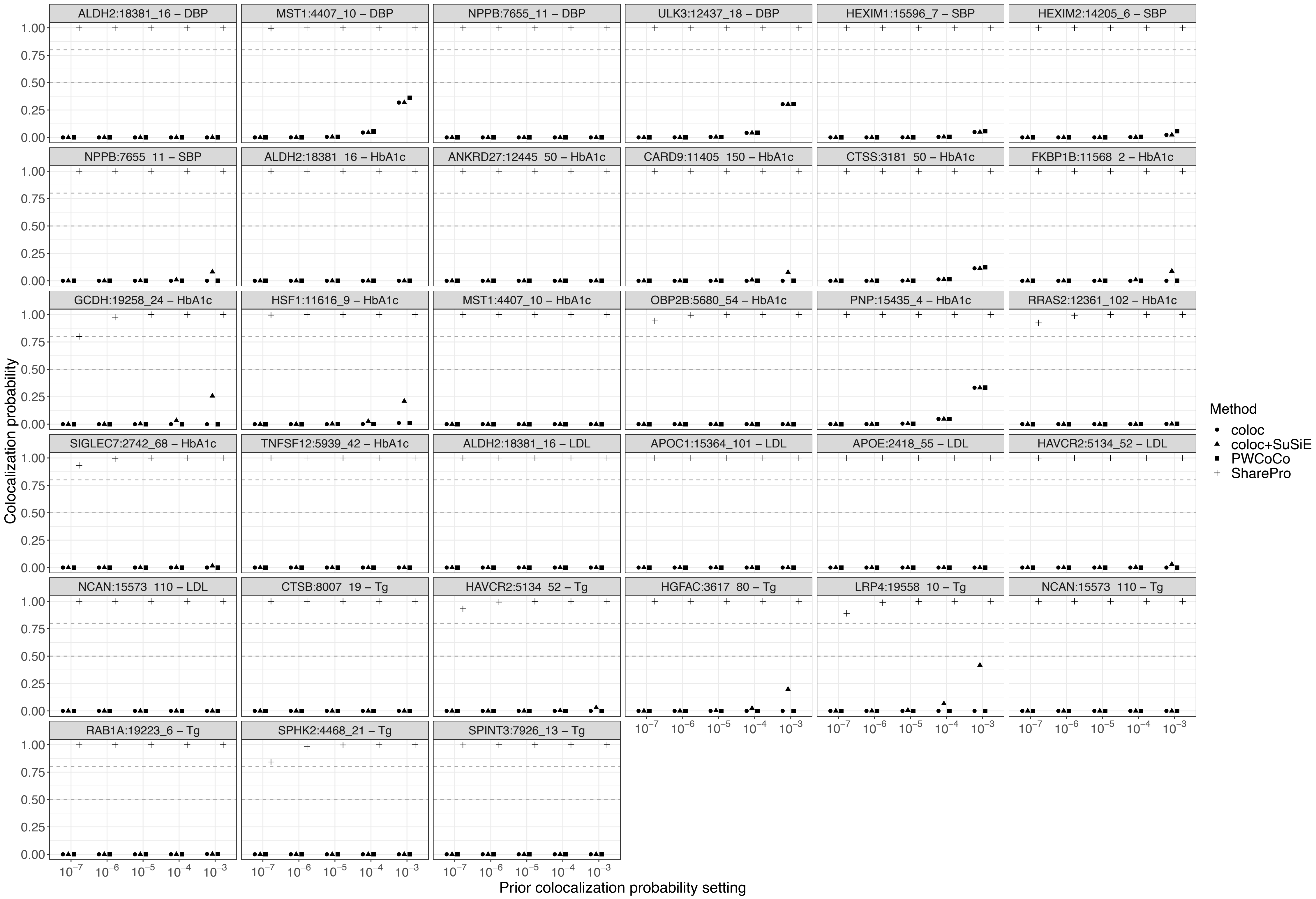


**Figure S19.** Prior sensitivity analyses for associations exclusively supported by SharePro. For each protein-trait association, colocalization probabilities inferred with different prior settings are indicated for each method. The grey dashed lines indicate colocalization probabilities of 0.8 and 0.5, respectively.
